# Adipose-mimetic granular hydrogels uncover biophysical cues driving breast cancer invasion

**DOI:** 10.1101/2025.10.23.684224

**Authors:** Brianna K. Knode, Garrett F. Beeghly, Brittany E. Schutrum, Dong Wang, Yitong Zheng, Alice Battistella, Ruchi Goswami, Chi-Yong Eom, Aline Bozec, Jochen Guck, Nozomi Nishimura, Salvatore Girardo, Corey S. O’Hern, Claudia Fischbach

## Abstract

1.

Breast cancer cells invade mammary adipose tissue during initial stages of metastasis but how the physical properties of adipose tissue regulate this process remains unclear. Here, we combined single cell mechanical characterization of primary adipocytes with microfluidic hydrogel fabrication, quantitative multiparametric imaging, Discrete Element Method (DEM) simulations, and *in vivo* experiments to elucidate these connections. First, we quantified the heterogeneous size and stiffness of primary adipocytes, and replicated these properties by fabricating adipocyte-sized polyacrylamide (PAAm) beads with tunable elasticity. Subsequently, we embedded these beads into type I collagen, the primary fibrillar extracellular matrix (ECM) component of breast adipose tissue, to form 3D granular hydrogels mimicking aspects of native adipose tissue architecture. Granular hydrogels embedded with beads demonstrated increased breast cancer cell invasion relative to bead-free controls, an effect that was more pronounced with soft versus stiff beads and correlated with increased collagen fiber alignment and hierarchical organization. In addition, live cell imaging and DEM simulations revealed that soft beads promoted invasion relative to stiff beads by deforming in response to confined cancer cell migration. Fiber alignment and adipocyte deformation trends were validated *in vivo* via intravital imaging of cancer cell migration in mammary fat pads of mice, and suggest that adipocyte mechanics regulate breast cancer invasion by coordinating both ECM architecture and cellular confinement. Ultimately, this work highlights the utility of tunable PAAm bead-collagen composites as micromechanical models to study the effect of adipose tissue structure on cancer cell invasion.

**Graphical Abstract:**
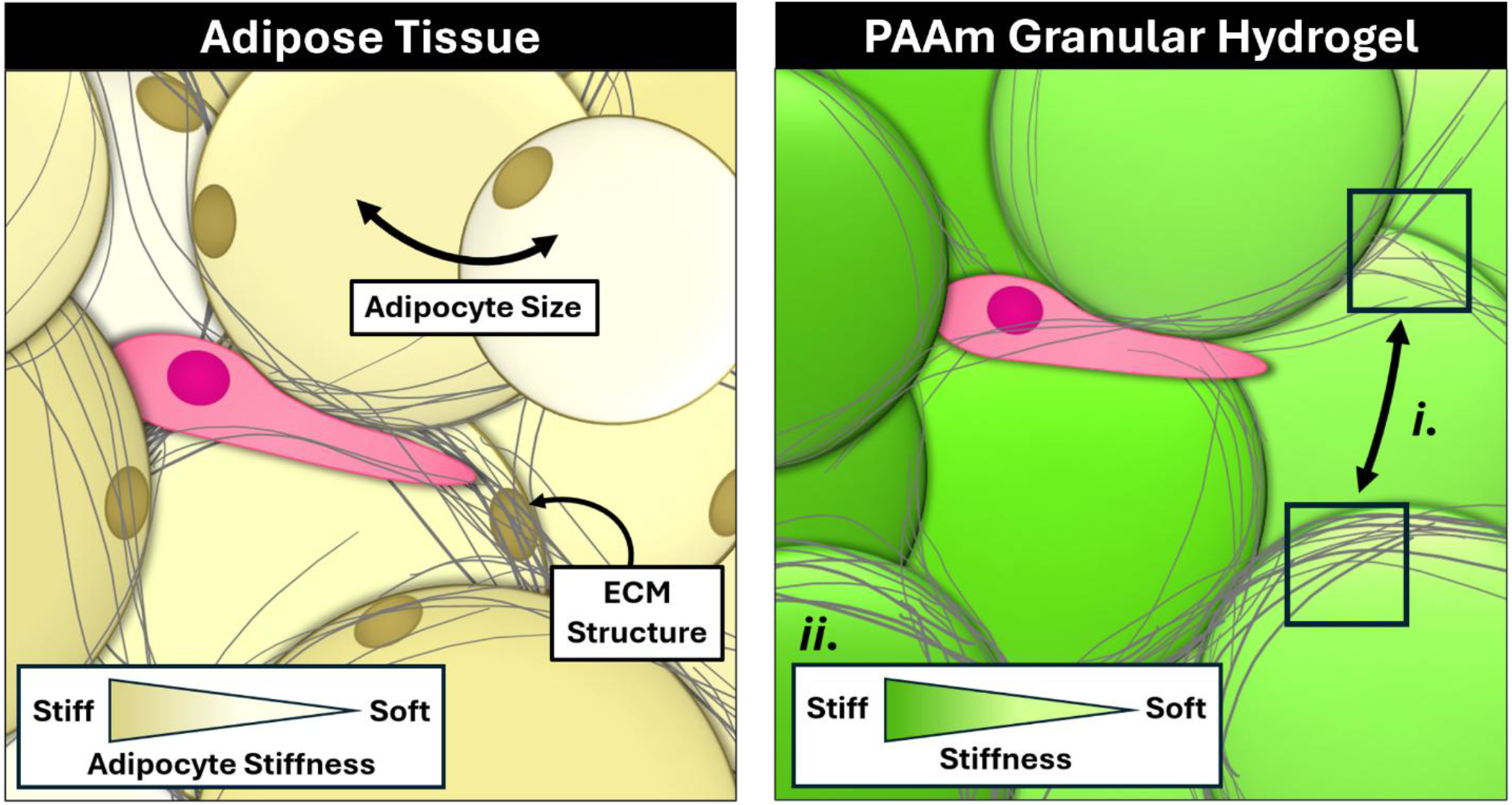
Polyacrylamide bead granular hydrogels recapitulate adipose tissue structure and alter cancer cell invasion in a stiffness-dependent manner. The invasion of primary breast cancer cells through mammary adipose is influenced by the heterogeneous mechanical properties of native stromal components, namely adipocytes and extracellular matrix ***(left*)**. Customizable granular hydrogels provide insight into the physical regulation of breast cancer progression by mimicking adipose tissue extracellular matrix structure (collagen I, *i*.) and stromal cell mechanics (polyacrylamide bead, *ii*.), whereby the inclusion of soft beads and/or dilute matrix promotes cancer cell migration compared to stiffer hydrogels ***(right*)**.

## 2. Introduction

Metastatic breast cancer is the leading cause of cancer-related death in women worldwide^1^. Despite decades of progress, our ability to diagnose and treat metastatic breast cancer remains limited due to an incomplete understanding of how cancer cells invade. Although most breast cancers originate in the mammary ducts and lobules^2^, metastasis begins when cancer cells escape these epithelial structures and migrate into the surrounding adipose tissue-rich host microenvironment^3,4^. They do so by squeezing between stromal structures, either as single cells or collective strands^5,6^, to reach blood vessels and ultimately colonize distant organs^7^. While previous studies have investigated how adipose tissue influences tumor cell phenotype through biochemical cues^8–10^, the role of its distinctive biophysical properties, which may be equally important, remains less clear.

About 85 percent of adipose tissue by volume is comprised of adipocytes, which are structurally and functionally unique due to their large size, spherical morphology, and lack of cell motility^11,12^. To support their roles in energy storage and thermoregulation adipocytes also exhibit physical and mechanical heterogeneity^13–15^. Notably, in metabolically altered states such as obesity and diabetes, adipocytes can adopt abnormal mechanical phenotypes^15,16^. Adding to this physical complexity, mammary adipocytes are surrounded by a network of extracellular matrix (ECM) primarily composed of fibrillar collagen type I^17^, which cancer cells navigate during invasion. In obesity, this collagen network becomes denser and more aligned^18,19^, which has been shown to promote tumor cell invasion in other contexts by influencing cell adhesion and physical confinement^19–22^. However, the relationship between adipocyte mechanics, ECM architecture, and breast cancer invasion remains poorly understood due, in part, to a lack of model systems capable of capturing the interplay between these different parameters in a tractable manner.

Most *in vitro* studies of adipose tissue employ one of three tissue culture methods, each with intrinsic limitations^23^. Most commonly, adipocytes are pre-differentiated from progenitor cells, but these cells do not develop the large and unilocular phenotypes characteristic of mature adipocytes and fail to recapitulate the mechanics of primary tissues^15^. In contrast, primary adipocytes exhibit the native structure and properties of the tissue from which they are resected, but are notoriously difficult to culture due to their buoyancy and fragility^24^. Moreover, isolated primary adipocytes^25,26^ and whole adipose tissue explants^27^ experience rapid phenotype loss, limiting their suitability for extended *in vitro* studies and manipulation of mechanical properties. Hence, no suitable methods currently exist to recapitulate the key mechanical and structural properties of native adipose tissue, motivating the need for innovative model systems to examine their effects on cancer cell invasion independently of biochemical signals.

While the physical characteristics of complex tissues are often modeled using homogenous bulk hydrogels^28^, granular hydrogels consisting of tunable microgel particles offer functional advantages^29–38^. For example, granular hydrogels demonstrate inherent porosity, control over hydrogel structure by adjusting particle packing density, and the ability to curate functionally diverse synthetic tissues via selective particle functionalization and interparticle crosslinking^29,30,36–39^. Artificially crosslinked packings of hydrogel particles are most commonly used to engineer tissues for regenerative approaches and study constituent cell dynamics^31–34,39^. In addition, some systems have combined hydrogel particles with ECM components^35,38^ or have tested the effect of cell packing or particle size on cancer growth^37,38^. However, how collagen network microarchitecture is controlled in such granular systems and the effects these changes have on invasion at varied bead stiffnesses have yet to be investigated. Consequentially, the fundamental relationships between adipocyte mechanics, fibrillar matrix architecture, and mammary cancer invasion remain unclear.

To model the discrete and continuous physical characteristics of adipose tissue, we designed adipocyte-sized hydrogel granules and embedded them into collagen type I at a physiologically relevant volume ratio. We first measured the effective diameter and stiffness of primary adipocytes and then recapitulated these parameters with adipocyte-mimetic polyacrylamide (PAAm) beads yielding biocompatible, cell-scale mechanical models with spherical morphology and mechanical tunability^40,41^. Moreover, incorporation of the beads into collagen I allowed us to replicate bulk adipose tissue organization and identify how adipocyte packing impacts collagen. Finally, we applied this biphasic model to experimentally and computationally investigate the role of both adipocyte and ECM mechanics on tumor cell migration, and validated our findings by investigating mammary tumor invasion in a syngeneic mouse model using intravital imaging. By mimicking adipose tissue’s biophysical traits, granular collagen hydrogels offer insights into how local stiffness and structure regulate cancer cell invasion. As adipose infiltration can influence cell function in various other contexts including metabolic diseases, our model provides a broadly applicable platform to study these connections.

## 3. Materials & Methods

### 3.1 Murine adipose tissue harvest and histology

For animal studies, 28,000 ES272 cells were injected into the ducts of the fourth mammary glands of female C57BL6 mice (The Jackson Laboratories) bilaterally. ES272 cells are a syngeneic C57BL6 breast cancer cell line with constitutively active and mutated PI3KCA (H1047R) (gifted from Dr. Ramon Parsons, Mount Sinai). Eleven days post injection, animals had reached humane endpoints. Animals were weighed and final tumor volumes were measured with calipers. Mice were euthanized via CO^2^ inhalation, then resected adipose tissue and mammary tumors were fixed in 4% (w/v) paraformaldehyde in 1x PBS for 18 hours at 4°C. Resected tumors were then stored in 70% (v/v) ethanol in water at 4°C until processing. Samples were sent to the Cornell College of Veterinary Medicine Animal Health Diagnostic Center for paraffin embedding, sectioning, and staining with hematoxylin and eosin or Masson’s trichrome. Stained sections were imaged and digitized on an Aperio ScanScope CS2 (Leica) with a 40x objective. All animal protocols were approved by the Institutional Animal Care and Use Committees at Weill Cornell Medicine and Cornell University.

### 3.2 Primary adipocyte isolation

Subcutaneous adipose tissue was collected from male C57BL6 mice (The Jackson Laboratories) at 14 weeks of age after 6 weeks of high fat diet (Research Diets, cat# D12330) fed ad libitum^42^. Primary adipocytes were then isolated from resected adipose tissue as previously described^43^. Briefly, adipose tissue was minced for several minutes until separated into approximately 1 mm^3^ pieces in Krebs-Ringer HEPES buffer (116 mM NaCl, 25 mM HEPES, 4 mM KCl, 2 mM D-glucose, 1.8 mM CaCl_2_, 1 mM MgCl_2_) supplemented with 1% (w/v) bovine serum albumin (KRHB). Floating tissue pieces were then digested in 1.5 mg/mL collagenase type I (Worthington Biochemical #LS004197) in Hank’s Buffered Salt Solution (Gibco #14065056) supplemented with 1% (w/v) bovine serum albumin for 50 minutes at 37°C. The resulting cell suspension was passed through 200 μm cell strainers (pluriSelect #435020003) to remove undigested pieces of tissue. Adipocytes were allowed to float out of solution for 10 minutes before the infranatant was aspirated and fresh KRHB was added. This process was repeated three times to enrich for adipocytes and remove contaminating stromal vascular cells.

### 3.3 Tunable polyacrylamide bead fabrication

Alexa Fluor™ 488-labeled polyacrylamide beads were produced and analyzed as previously described^40,41^. Briefly, a polydimethylsiloxane (PDMS)-based microfluidic chip employing a flow-focusing geometry was utilized to produce polyacrylamide pre-gel droplets. The chip design incorporates two inlets for the polyacrylamide pre-gel mixture and oil flows, and one outlet for droplet collection. The chip features a cross-junction with a width of 40 μm, which gradually widens downstream to 50 μm, 100 μm, and 150 μm, while maintaining a constant channel height of 60 μm.

The polyacrylamide pre-gel droplets were produced in fluorinated oil (3M™ Novec™ 7500, Iolitec Ionic Liquids Technologies GmbH) containing ammonium Krytox^®^ surfactant (2.4%w/v) as an emulsion stabilizer, N, N, N′,N′ -tetramethylethylenediamine (TEMED) (0.4%v/v) (Sigma-Aldrich) as a catalyst and acrylic acid N-hydroxysuccinimide ester (NHS) (0.1%w/v) (Sigma-Aldrich) to include NHS functional groups into the final gel meshwork for binding of Alexa Fluor™ 488 hydrazide. The pre-gel mixture contained acrylamide (40%w/v) (Sigma-Aldrich) as a monomer, bis-acrylamide (2%w/v) (Sigma-Aldrich) as a crosslinker, ammonium persulphate (0.05%w/v) (Sigma-Aldrich) as a radical initiator and Alexa Fluor™ 488 hydrazide (2 mg/ml) (Thermo Fisher) diluted in 10mM Tris buffer (pH = 7.48). Total monomer concentration in the pre-gel mixture together with the droplet diameter was fine-tuned to obtain beads with well-defined diameter and elasticity. After in-drop polymerization, the beads were washed and resuspended in 1xPBS (pH = 7.4). The diameter of the beads in PBS was analyzed by acquiring bright-field images of the beads (Zeiss AxioObserver.A1, A-Plan 10x/0.25 objective) and by analyzing them using a macro implemented on an open-source Fiji. Elasticity was measured by Atomic Force Microscopy (AFM) indentation.

### 3.4 Atomic force microscopy

Isolated adipocytes were immobilized to glass-bottom Petri dishes (World Precision Instruments #FD35100) pre-coated with 5 μg/cm^2^ Cell-Tak Cell and Tissue Adhesive (Corning #354240) per manufacturer’s instructions. In short, freshly isolated primary adipocytes were tightly packed by aspirating infranatant buffer and supernatant free lipid as previously described^43^, then pipetted directly onto the prepared Cell-Tak substrate. Adipocytes were maintained in a 37°C incubation chamber and allowed to attach for several minutes before being submerged in imaging media. A subset of sample was stained with Hoechst prior to AFM to confirm that the isolated structures were true adipocytes, rather than residual lipid droplets (**Fig. S1b**). For AFM indentation experiments on adipocytes and PAA beads, a Nanowizard IV was used (JPK BioAFM, Bruker Nano GmbH). PNP-TR-TL-Au cantilevers (NanoWorld) with a nominal spring constant of 0.32 N/m were employed. These probes have been previously modified with a polystyrene bead of 5 μm diameter (microparticles GmbH) and calibrated by the thermal noise method on the surface of a glass slide with milli-Q water. The measurements were performed directly in the Petri dishes on the surface of which adipocytes or PAA beads were immobilized as described above. A maximum force load of 15 nN at a rate of 5 μm/s in z closed-loop feedback mode was used. AFM force-indentation analysis was performed using the open-source analysis software PyJibe (https://github.com/afm-analysis/pyjibe; version 0.14.0). The Poisson’s ratio was set to 0.5 for all analyses. We applied the Hertz/Sneddon model for a spherical indenter with a limitation of the indentation curve to 1.5 μm from the contact point. A geometrical correction factor (*k*) was applied to correct the indentation for the additional deformation of the spherical shape coming from the substrate (Glaubitz et al., *Soft Matter*, 2014). The correction factor *k* was calculated using

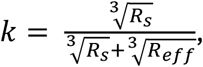

where *R*_*s*_ is the sample radius, and the effective radius (*R*_*eff*_) is given by

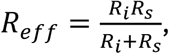

where *R*_*i*_ is the indenter radius. Further details of this procedure can be found in the documentation of the Phyton Library for Nanite (https://nanite.readthedocs.io/en/stable/sec_fitting_guide.html#geometrical-correction-factor).

### 3.5 Second harmonic generation imaging

Multiphoton imaging was performed on unstained adipose tissue sections with an inverted Zeiss 880 confocal/multiphoton laser scanning microscope. Imaging was performed with non-descanned detection, 780 nm excitation, and an Olympus 10x/0.45W Apochromat objective. Emitted light was filtered for second harmonic generation (360–405 nm) to visualize collagen fibers.

### 3.6 Microwell fabrication

Customizable PDMS microwells were constructed using the Sylgard 184 Silicone Elastomer Kit (Dow #000000818156). Reagents A (polymer base) and B (curing agent) were combined at a 10:1 ratio respectively, and vigorously mixed for 5 minutes. This mixture was transferred to a 150mm diameter polystyrene Petri dish, degassed for 10 minutes in a vacuum chamber, and cured overnight at 60°C. Once cooled, microwells were cut from the cured PDMS using Integra Miltex 4mm (inner well diameter), 8mm (outer well diameter), and 10mm (lid) disposable biopsy punches, and stored in a 100mm diameter Petri dish. The PDMS lids were submerged in 70% (v/v) ethanol in water, vigorously shaken on a rocker for 20 minutes, and allowed to air dry under sterile conditions. Single 18mm round #1 cover glasses (VWR #16004-300) were similarly washed and dried, then placed in the wells of a 12-well cell culture plate (Falcon #351143). The 12-well plate and microwell-containing Petri dish were plasma treated (Harrick Plasma) on the highest setting for 1 minute. Microwells were then transferred to the 12-well plate and bonded to the glass coverslips via an additional 5-minute round of air plasma treatment. For live cell tracking experiments, PDMS microwells were bonded to glass-bottomed culture dishes (VWR #10810-054) rather than circular coverslips. Microwell surfaces were lastly treated with 1% (v/v) poly-ethylenimine (PEI, Sigma-Aldrich #181978) for 10 minutes, 0.1% (v/v) glutaraldehyde (Sigma-Aldrich #G7651) for 30 minutes, thoroughly rinsed with sterile water and stored at 4°C.

### 3.7 Hydrogel preparation

To fabricate the hydrogels, collagen-bead mixtures were prepared over ice and cast into chilled PDMS microwells. Rat tail collagen I (Corning #354249) was diluted to a final concentration of 2.5 mg/mL or 6 mg/mL in high glucose DMEM cell culture medium (Gibco #12800082) and neutralized with 1N NaOH. PAAm bead stocks of varying stiffness were briefly spun down in a mini centrifuge and the 1x PBS supernatant was aspirated before gently resuspending the beads in one of the prepared collagen solutions at a 6:1 ratio, respectively. Collagen solutions with and without beads were pipetted into PDMS microwells, such that the wells were slightly overfilled, and PDMS lids were oriented on top of the filled wells to avoid air bubbles. Hydrogels were polymerized via the cold-cast protocol as previously described^44^; in short, gels were allowed to incubate for 15 minutes at 4°C, room temperature, and 37°C, and were flipped every 5 minutes during the polymerization process to ensure bead packing consistency throughout the gel. After a total of 45 minutes, the hydrogels were submerged in the high glucose DMEM, PDMS lids were removed, and gels were stored at 37°C for 36 hours. The hydrogels were finally fixed in 4% paraformaldehyde (PFA) for 25 minutes, then thoroughly washed and stored in 1x PBS at 4°C.

### 3.8 Bicinchoninic acid assay for collagen quantification

The collagen concentrations of pregel solutions were quantified using a micro-BCA protein assay (Thermo Scientific #23235). Experiments were conducted according to manufacturer specifications, with the following exceptions. Rat tail collagen type I solutions were serially diluted in milliQ water to eight final concentrations of 80 μg/mL, 40 μg/mL, 20 μg/mL, 10 μg/mL, 5 μg/mL, 2.5 μg/mL, 1 μg/mL, and 0.5 μg/mL, and were used to establish experimental standard curves. Beadless collagen solutions (2.5 mg/mL or 6 mg/mL) were prepared as previously described, and 1:6 collagen-to-bead mixtures were prepared from these stocks. These beadless and bead-laden collagen solutions were diluted in milliQ water to a target total collagen concentration of 40 μg/mL, excluding the bead volume fraction from the calculations. PAAm beads were allowed to settle out of the solutions during the colorimetric reaction, such that only the reacted collagen in the supernatant was analyzed. All absorbance measurements were acquired within 10 minutes of each other using a Beckman Coulter DU730 Life Science UV/Vis Spectrophotometer.

### 3.9 Cell culture

MDA-MB-231 (ATCC) and EO771-YFP (LanYFP-expressing, CH3-Biosystems, provided by the Schaffer-Nishimura lab at Cornell University) cells were cultured in 75cm^2^ flasks using high glucose DMEM cell culture medium (Gibco #12800082) supplemented with 10% fetal bovine serum (FBS, Atlanta Biologicals #S11150) and 1% penicillin/streptomycin (Gibco #15070063). MCF10A cells (ATCC) were similarly cultured using high glucose DMEM/F12 culture medium supplemented with 5% horse serum (Thermo Scientific #16050122), 10 μg/mL insulin (Krackeler Scientific #45-91077C), 500 ng/mL hydrocortisone (Thermo Scientific Chemicals #A16292-03), 100 ng/mL cholera toxin (Krackeler Scientific #45-C8052), 20 ng/mL EGF (Invitrogen #PHG0313), and 1% penicillin/streptomycin. Cells were incubated at 37°C in an atmosphere of 5% CO_2_ and maintained for experimental use until passage 20.

### 3.10 Invasion and migration assays

Breast cancer cells were suspended in the appropriate high glucose DMEM cell culture medium at a concentration of approximately 133,333 cells/mL. For endpoint experiments, polymerized hydrogels were submerged in 1.5mL of this cell suspension and incubated at 37°C/5% CO_2_ for 36 hours. For live cell tracking, the cell suspension was first incubated with 100ng/mL Hoechst stain (Thermo Scientific #H3570) for 45 minutes at 37°C, then centrifuged and resuspended in fresh media. Hydrogels cast in glass-bottomed culture dishes were submerged in 2mL of this cell suspension and incubated at 37°C/5% CO_2_ for 21 hours during spinning disk confocal image acquisition. After incubation, all hydrogels were removed from cell suspension, fixed in 4% PFA solution for 25 minutes, then thoroughly washed and stored in 1x PBS at 4°C.

### 3.11 Immunofluorescence

Fixed hydrogels were submerged in a permeabilization solution comprised of 0.1% Triton X-100 (Thermo Scientific #AAA16046AP) and 1% bovine serum albumin (BSA, Fisher Scientific #BP1600-100) in 1x PBS for 15 minutes at room temperature. Permeabilization solution was aspirated and a 132nM solution of Alexa Fluor 568-conjugated phalloidin (Thermo Scientific #A12380) in 1x PBS + 1% BSA was pipetted on top of the exposed hydrogel surface in a small-volume bleb. The staining solution was allowed to incubate for 45 minutes at room temperature. This process was repeated a second time to ensure proper diffusion of the staining solution throughout the gel. The hydrogels were lastly submerged in a 715nM solution of DAPI (Thermo Scientific #D1306) in 1x PBS + 1% BSA for 30 minutes at room temperature before being rinsed and stored in 1x PBS at 4°C.

### 3.12 Confocal microscopy

For 36-hour endpoint experiments, PDMS ring-encapsulated hydrogels were removed from the 12-well plate, inverted, and mounted in 1x PBS on a 24x60mm No.1 glass coverslip (VWR #48404-455). Representative image stacks were acquired on a Zeiss LSM880 confocal multiphoton inverted microscope using a C-Achroplan 32x/0.85 W Corr M27 objective. Z-stack images of the top 150μm of each gel were acquired with step sizes of 4μm or 5μm, each slice measuring 512 pixels x 512 pixels. The Zeiss Zen Smart Setup function was used to configure fluorescence channels for DAPI, Alexa Fluor 488, and Alexa Fluor 568 visualization. Reflectance imaging was used to evaluate collagen fiber structures, for which a customized reflectance channel was added to the Smart Setup configuration.

For live cell tracking experiments, time-lapse image stacks were acquired on an Olympus IX83 inverted spinning disk confocal microscope using a 10x/0.4 air objective. Two fluorescence channels were configured to visualize DAPI and Alexa Fluor 488 signal. Z-stack images of the top 100μm of the gels (5μm step size) were acquired every 15 minutes over 21 hours, and three representative fields of view were collected for the hydrogel imaged.

### 3.13 Mammary tumor imaging window implantation

Three 6-month-old female C57BL/6 mice (The Jackson Laboratory) were anesthetized using isoflurane inhalation. LanYFP-expressing EO771 mammary carcinoma cells (1 × 10^4^ cells in 20 μL PBS) were orthotopically injected into the third mammary fat pad under aseptic conditions. Tumor growth was monitored by palpation. Two weeks post-injection, when palpable tumors were established, animals underwent surgical implantation of a mammary imaging window. Briefly, after hair removal and disinfection, a circular titanium frame (12 mm inner diameter; APJ Trading Co.) was positioned over the tumor region and secured to the skin using interrupted sutures. A sterile glass coverslip (No. 1 thickness) was affixed over the frame to enable repeated optical access to the tumor microenvironment as described previously^45^.

### 3.14 Intravital two-photon microscopy

Intravital imaging was performed using a multiphoton microscope equipped with a Chameleon Ti:sapphire laser (Coherent) tuned to 830 nm for excitation as published previously^46^. Fluorescence was collected in three detection channels of the microscope formed by three long-pass dichroics: LM01-488, FF560-Di02, FF593-Di03. In addition, the channels had the following bandpass filters respectively, FF01-417/60, FF01-517/65, and FF01-629/53. SHG signal was collected for collagen fibers, and fluorescence signal was collected for LanYFP tumor cells and Texas Red–dextran-labeled blood plasma (70 kDa; Thermo Fisher Scientific, 30 μL of 2.5% in saline injected intravenously). Imaging was conducted with a 20× water-immersion objective at 37°C. Two to three representative regions of interest measuring at least 216 μm x 216 μm x 100 μm (512 pixels x 512 pixels x 100 voxels) were acquired for each mouse across several timepoints. Care was taken to maintain anesthesia throughout imaging in accordance with Cornell University IACUC protocol.

### 3.15 Image analysis

All acquired images were processed and analyzed using the open-source image analysis software ImageJ/Fiji^47^, unless otherwise noted. Sample counts for each analysis are included in the respective subsection, and comprise at least 3 biological replicates unless stated differently.

#### Adipocyte/PAAm bead characterization

To evaluate the distribution of adipocyte diameters in murine adipose tissue, histological sections stained with Masson’s trichrome were evaluated. The areas of individual adipocyte cross-sections were evaluated, and undeformed adipocyte diameters *d* were calculated using 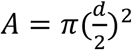 where *A* is the adipocyte cross-sectional area. 1626 individual adipocyte cross-sections were analyzed, collected from 5 mice. Similarly, to analyze the diameter distributions of PAAm bead batches, brightfield images of PAAm beads were acquired and manually traced in ImageJ/Fiji. Since PAAm beads maintain high sphericity after polymerization, the diameters of individual beads were directly traced rather than using the circumference to infer an approximate diameter.

Confocal image stacks were utilized to analyze the local packing fraction of PAAm beads within each collagen hydrogel condition via the tracking of bead centers and diameters; one z-stack was acquired for each condition, and each stack depicted approximately 6 layers of bead packings. First, a circular Hough transform was used in MATLAB to identify the circular cross-sections along with their centers and radii in each image slice. We denote the x-y plane to be parallel to the image slice, and the z-direction parallel to the confocal microscopy scanning direction. Second, we identify whether circular cross-sections from different z-slices belong to the same bead. Any circles whose centers are no more than 0.1 of the bead diameter apart along the x-y plane and no more than the bead diameter apart along the z-direction are deemed to belong to the same bead. Third, using the x, y, and z positions of all circular cross-sections deemed to be from the same bead, we obtain the center and radius of that bead through least-square fitting to the equation for a sphere. After determining the centers and radii of the beads, we further calculate the local packing fraction for each tracked bead. To do this, we first apply a Voronoi tessellation to the bead packing using Voro++^48^. We then calculate the local packing fraction defined as the bead volume divided by the volume of its associated Voronoi polyhedron.

#### Collagen fiber network analyses

Granular hydrogel collagen fiber characteristics were analyzed using multiple computational platforms; 150-μm image stacks collected from 5 regions of interest (ROIs) in each condition were first corrected to eliminate noise. Confocal reflectance microscopy visualization of these collagen hydrogel networks yielded raw micrographs with high background signal and a central artifact that interfered with downstream analyses. To minimize the contaminating noise, all raw reflectance images were subjected to the ImageJ/Fiji “Subtract Background” function at a sigma value of 5 pixels. To eliminate the central reflectance artifact, the ImageJ/Fiji “Image Calculator” was used to identify the artifact via overlap of multiple image slices (“AND” function) and then subtract the isolated artifact from each raw image slice (“SUBTRACT” function), after which image stacks were automatically thresholded using the “Triangle” filter.

The collagen fiber networks in living murine adipose tissue were similarly analyzed using SHG images collected from intravital two-photon microscopy of mammary tumors. SHG image stacks (100 μm x 100 μm x 40 μm (xyz) with a step size of 1 μm) were collected from two mice (two ROIs of the same tumor per mouse). These four image stacks were used for analysis, each representative of a different tissue structure: highly fibrotic or less fibrotic, with or without packed adipocytes (adipocyte identification details below). To address the high background signal, which is characteristic of intravital imaging, and better illuminate the collagen structures to be characterized, the ImageJ/Fiji “Subtract Background” function was applied at a sigma value of 10 pixels (4.22 μm). Image stacks were then automatically thresholded using the “IsoData” filter. To quantify the differences in fibrosis between representative fields of view, the ImageJ/Fiji “Histogram” function was used to calculate the number of positive (above threshold) and negative (background) pixels for each slice. The average ratio of these values was computed for each representative stack, and compared between conditions such that a larger positive pixel fraction was indicative of increased fibrosis.

The alignment of collagen fibers both *in vitro* and *in vivo* were evaluated using the ImageJ/Fiji plugin OrientationJ “Analysis” function, as has been previously reported^49^. In short, fiber “coherency” defines the relative orientation of locally defined image features by employing a gradient structure tensor that accounts for relevant directional information, whereby a coherency score of 1 is suggestive of perfect anisotropy at the defined scale and a score of 0 indicates local fiber isotropy. For this specific protocol, a local window size of 6 pixels, approximating the size of a focal adhesion complex^50^, was used to survey confocal reflectance microscopy images of collagen fibers via the cubic spline method. Image output parameters were adjusted such that the hue of the collagen network corresponded to the local fiber coherency score. The RGB images resulting from this color survey were converted to HSB stacks, from which the Hue channel was isolated and 8-bit histogram values (0-255) were collected. 8-bit values were converted to a 0-1 scoring scale corresponding to fiber coherency scores, and score distributions were evaluated for outliers using the GraphPad Prism ROUT method. The maximum desired False Discovery Rate (FDR), denoted by the variable Q, was set equal to 1%. Identified outliers were removed, and cleaned datasets were analyzed to draw conclusions about local collagen fiber alignment.

Granular hydrogel collagen fiber interconnectivity was observed and characterized by thresholding the aforementioned Hue channel of each color-surveyed image at intervals corresponding to coherency scores from 0-1. The “3D Object Counter” function was then used to identify fiber clusters greater than 250μm^3^ in volume such that any remaining reflectance artifacts and/or single collagen fibers were removed from the binarized output image. Each binary 3D object identified a distinct cluster of interconnected fibers, and the volumes of these objects were collected at each coherency score threshold. The overall matrix connectivity score at each threshold level was defined as follows:

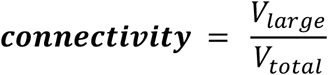

where *V*_*large*_ is the total volume of all identified large objects (large, interconnected fibers) and *V*_*total*_is the total volume of all identified objects (both small, single fibers and large, interconnected fibers) after coherency score thresholding. The connectivity score measures the volume fraction of large, interconnected collagen fibers among all collagen fibers. The connectivity score ranges between 0 and 1 (or 0 and 100, depends on whether we want to use percentage), with 0 indicating no presence of large fibers and 1 indicating all identified collagen fibers are interconnected. Connectivity scores were calculated for each coherency score threshold level and compared between mechanically distinct hydrogel conditions.

To further analyze the local alignment of the collagen fibers^51^, a subset of confocal images (one z-stack, 31 slices per condition) of collagen was divided into small, overlapping 20.75 μm by 20.75 μm windows (or 40 pixels by 40 pixels, with each two adjacent windows overlapping by 50%). Then, each window was transformed into a window with the same size F in the Fourier space using a 2D fast Fourier Transform (FFT). The central image moments of the FFT were calculated as follows:

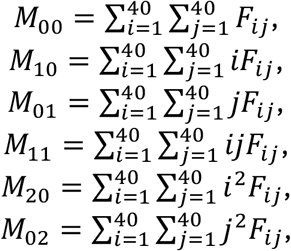

where i and j are indices of a pixel along the horizontal and vertical directions, respectively, and F_ij_ is the value of the Fourier transformed window at pixel (i, j). The central moments were then used to construct the 2 by 2 symmetric covariance matrix of the window image:

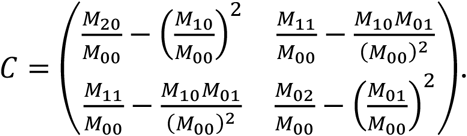

The direction of the eigenvector corresponding to the larger eigenvalue of C reveals the direction of collagen fibers in that window, θ, and the eigenvalues (λ_1_ and λ_2_) of C are used to calculate the eccentricity of collagen fiber alignment, e=| λ_1_ -λ_2_|/(λ_1_ + λ_2_). The eccentricity ranges between 0 and 1, with 0 corresponding to random collagen orientations and 1 corresponding to perfect collagen fiber alignment. Windows with low enough image intensity (10% of the highest image intensity) were ignored for this analysis.

Lastly, the same subset of confocal image slices (one z-stack, 31 slices per condition) was used to evaluate the structural characteristics of individual collagen fibers using the independent MATLAB application CurveAlign^52^. To measure the approximate length, width, and straightness of individual collagen fibers, corrected reflectance images of collagen networks were opened in the CT-Fire function of the CurveAlign application. CT-Fire parameters were configured as follows: ‘thresh_im2’ = 15, ‘s_xlinkbox’ = 5, minimum fiber length = 4 pixels, fiber line width = 0.5, maximum fiber width = 50 pixels, and the percentile of remaining curvelet coefficients was set to 0.2. Output fiber length, width, and straightness distributions were averaged and compared between the mechanically distinct hydrogel conditions.

#### Cancer cell invasion analyses

MDA-MB-231 cell invasion through control and bead-laden collagen hydrogels was investigated by tracking individual cell nuclei in the z-direction after 36 hours of invasion. For each mechanical condition, 4 confocal image stacks were analyzed, and DAPI-labeled nuclei were identified using the TrackMate plugin configured with the StarDist tracker for segmentation^53,54^. All TrackMate parameters remained set to the default values with the exception of the following adjustments: auto quality and contrast filters were applied, a simple LAP tracker was used to track individual nuclei, the linking max distance was set to 5 pixels, the gap-closing max distance to 5 pixels, and the gap-closing max frame gap to 1 frame interval. Display tracks were excluded, and a label image including spots corresponding to tracks was exported for further analysis. The Fiji 3D Objects analysis tool was then used to evaluate the localization of individual nuclei throughout the acquired z-stack, and the centroid measurement of any object greater than 100 voxels in volume was recorded. The invasion depth of cells was calculated after a 36-hour incubation period as follows:

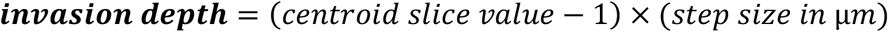

Additionally, the diffusion constant was calculated for the same populations of cells. If we assume that the cancer cells move diffusively, for each of the six mechanical conditions, the probability distribution of the invasion depth *z* follows a half-Gaussian distribution^55^:

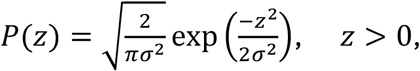

where 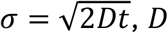 is the diffusion constant of the cancer cells, and *t* is the time elapsed since the beginning of the experiment. The cumulative distribution for a half-Gaussian process is

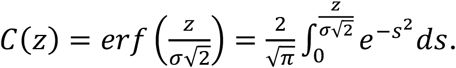

The experimentally measured mean of the invasion depth 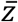 will converge to 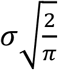 as the number of experimental measurements increases. Thus, we can calculate 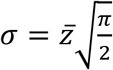 and the resulting diffusion constant

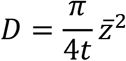

from the experimental measurements of the invasion. The cumulative distribution of the experimentally measured invasion depths and from an ideal half-Gaussian process with the best-fit values of *σ* (and corresponding R^2^ values) are shown in **Fig. S2d-e**.

Confocal micrographs of MDA-MB-231 cell nuclei squeezing between collagen-embedded PAAm beads were evaluated computationally to identify differences in nuclear deformation, and the shapes of the deformed nuclei were compared to those of unconfined nuclei. First, 15 confined nuclei were isolated from the segmented image stacks of each of the four bead-laden hydrogel conditions; nuclei selected for this analysis were in contact with two or more PAAm beads in the same x-y plane. Similarly, 15 nuclei uncontacted by PAAm beads during image acquisition were isolated from the same segmented image stacks. The nuclei of cells invading through beadless collagen hydrogels were also isolated for comparison. Lastly, the dimensionality of each isolated nucleus was reduced via maximum intensity projection, and 2D nuclear roundness was calculated by applying the Fiji “Measure” function to each replicate image.

A similar method was employed to characterize the deformation of PAAm beads in contact with migratory cancer cells. Confocal image stacks of PAAm bead hydrogels including and excluding cancer cells were surveyed, and at least 60 beads were manually segmented for each condition. Undeformed bead metrics were collected from image stacks of hydrogels not seeded with MDA-MB-231 cells, while deformed beads were defined as those in contact with and contributing to the confinement of migratory cell nuclei. Individual bead roundness was evaluated for all conditions by isolating the x-y plane at which the bead of interest was most visually deformed, then characterizing this segmented slice using the Fiji “Measure” function.

The same technique was applied to characterize the deformation of adipocytes *in vivo*. Briefly, intravital SHG (collagen) and fluorescent (separate EO771 and blood plasma) images were overlaid, and adipocyte boundaries were manually traced from a representative stack. The fluorescent dextran intended for vasculature labeling also diffused into surrounding tissue structures, presumably due to leaky tumor vasculature, and provided good contrast for adipocyte tracing. Adipocytes were identified as any relatively spherical volume of 3D void spaces greater than 20 μm in diameter. Adipocyte cross sections were manually segmented slice by slice, and regions of high and low cancer cell invasion were identified. Overlaying this masked stack with the corresponding EO771 image stack, adipocyte cross sections which were in contact with confined cancer cells were isolated for analysis. Adipocyte cross sections that were not in contact with confined cancer cells served as undeformed controls. Five representative cell masks for each sample type were evaluated using the Fiji “Measure” function to determine roundness and circularity values.

To characterize live cell invasion data acquired via spinning disk confocal microscopy, it was first necessary to track cancer cell nuclei localization over the time course. We identified cell nuclei centers at each time step. To do so, at a given time step, we binarized images taken from spinning disk microscopy at all z-stacks and then used “regionprops3” function in MATLAB to identify voxels that belonged to each cell nucleus. We then calculated the center of mass for each nucleus as the center for that nucleus. After identifying all cell nuclei centers from all time steps, we implemented the method described by Crocker and Grier^56^ to track the cells. We calculated the speed of a cell at a given time step as the displacement of that cell from the previous time step to the current time step divided by the time interval.

Furthermore, the relationship between the bead packing fraction and cell acceleration was evaluated by calculating the distance between a migratory cell’s nuclear centroid and the surface of nearby beads tracked from the spinning disk image stacks. It was first necessary to determine the precise packing of PAAm beads throughout the acquired region of interest. Z-stacks were surveyed manually whereby the largest x-y spherical cross section of each bead was segmented and duplicated into a temporary stack, maintaining the dimensionality of the original images. These segmented stacks were then uploaded to the 3D modeling software Dragonfly (license #BNR90-30900-G1HJK-M8P9Q-911AH-KPYED-GNU89KDUV) and the ‘Create a Sphere’ Shape Tool was used to reconstruct the volume of each bead throughout the stack. This was achieved by aligning the radius and center of a sphere with that of a segmented bead cross section, then replacing image stack data contained within the spherical region with saturated pixels using the ‘Image Operations’ Overwrite function. Once all bead volumes were reconstructed, the resulting stack was exported then merged with the original segmented nuclear channel in Fiji, and the localization of migratory nuclei between PAAm beads was quantified for every timepoint. Using the centers and radii of tracked beads, we calculated the distance between the centroid of a nucleus of interest and its nearest neighboring bead. Finally, the cell accelerations were determined from the migratory speed datasets, and then correlated with the degree of local confinement (bead proximity) calculated for each timepoint.

### 3.16 Discrete Element Method (DEM) modeling of breast cancer and adipocytes

Adipocytes are modeled as deformable particles and breast cancer cells as adhesive soft disks in two dimensions. The total energy of the system is given by

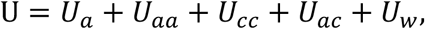

where *U*_*a*_ is the total shape energy of the adipocytes, *U*_*aa*_ is the total interaction energy between adipocytes, *U*_*cc*_ is the total interaction energy between cancer cells, *U*_*ac*_ is the total interaction energy between adipocytes and cancer cells, and *U*_*w*_ is the total interaction energy between both adipocytes and cancer cells and the confining walls. Each adipocyte is modeled as a deformable polygon with *N*_*v*_ vertices, and *U*_*a*_ is given by

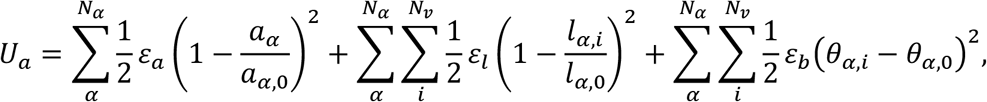

where *N*_*α*_ is the total number of adipocytes, *a*_*α*_ is the area of adipocyte *α* with a preferred area *a*_*α*,0_, *l*_*α,i*_ is the length between vertices i and (i+1) with a preferred length *l*_*α*,0_, and *θ*_*α,i*_ is the angle formed by vertices (i-1), i, and (i+1) with a preferred angle *θ*_*α*,0_. *ε*_*a*_, *ε*_*l*_, and *ε*_*b*_ are the energy scales that control deviations of adipocyte area, perimeter, and local curvature from their rest values, respectively. (See **Fig. 5a**.)

To prevent adipocytes from inter-penetrating, we include an energy penalty when two adipocytes overlap:

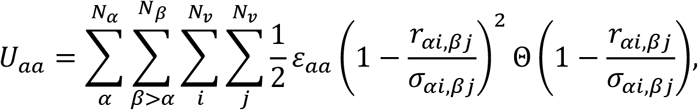

where *r*_*αi,βj*_ is the distance between vertex i on adipocyte *α* and vertex j on adipocyte *β, σ*_*αi,βj*_ is the sum of the radii of vertex i on adipocyte *α* and vertex j on adipocyte *β, ε*_*aa*_ is the repulsive energy scale, and Θ(·) is the Heaviside step function. (See **Fig. 5a**.) *N*_*c*_ cancer cells interact pairwise via the following energy:

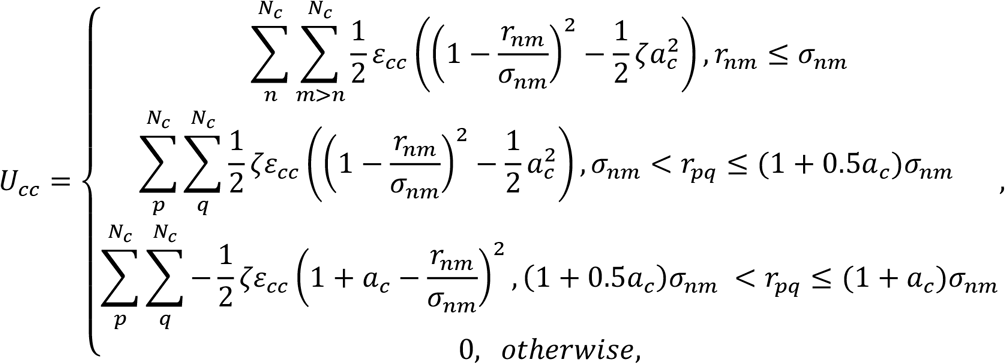

where *r*_*nm*_ is the distance between cancer cells n and m, *σ*_*nm*_ is the sum of the radii of cancer cells n and m, *a*_*c*_ sets the attraction range, *ε*_*cc*_ is the energy scale, and *ζ* sets the attraction strength. The resultant force is shown in **Fig. 5a**. Adipocytes and cancer cells interact via purely repulsive forces given by the gradient of the following potential energy:

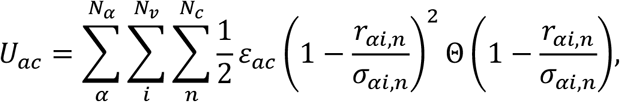

where *r*_*αi,n*_ is the distance between vertex i on adipocyte *α* and cancer cell n, and *σ*_*αi,n*_ is the sum of radii of vertex i on adipocyte *α* and cancer cell n. Packings of adipocytes and cancer cells are confined within a rectangular box with area *L*_*x*_*L*_*y*_ and periodic boundary conditions in the y-direction and confined by two walls in the x-direction. (See **Fig. 5a**.) One wall is fixed at *x* = 0, and the other is positioned at *x* = *L*_*x*_. Both adipocytes and cancer cells interact with the walls via purely repulsive forces that are generated via the gradient of the following potential energy:

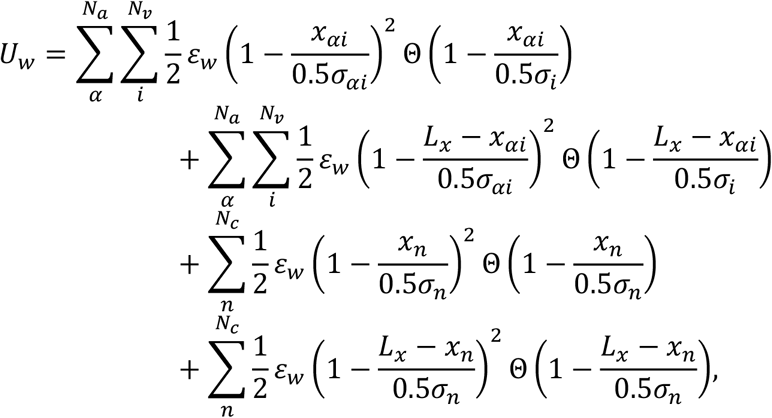

where *x*_*αi*_ and *x*_*n*_ are the x-positions of vertex i on adipocyte *α* and cancer cell n, respectively, *σ*_*αi*_ and *σ*_*n*_ are the diameters of vertex i on adipocyte *α* and cancer cell n, respectively, and *ε*_*w*_ is the energy scale of the interactions with the wall. The pressure *P* in the system is given by the sum of all forces exerted on the wall at *L*_*x*_, divided by *L*_*y*_.

We study systems with *N*_*α*_ = 32 adipocytes with half possessing *N*_*v*_ = 20 vertices and the other half possessing *N*_*v*_ = 28 vertices, and *N*_*c*_ = 800 cancer cells. For each adipocyte, we set the shape parameter 𝒜 = (*N*_*v*_*l*_*α*,0_)^2^ /(4*πa*_*α*,0_) = 1 and *θ*_*α*,0_ = 2*π*/*N*_*v*_, such that a stress-free adipocyte possesses the shape of a regular polygon with *N*_*v*_ sides. We set *a*_*α*,0_ = 5000 *μm*^2^ (10000 *μm*^2^) for the small (large) adipocytes with *N*_*v*_ = 20 (28) so that the average adipocyte diameter is ⟨*D*_*a*_⟩ = 100 *μm*. We draw the diameters of cancer cells from a uniform distribution with mean ⟨*σ*_*n*_⟩ =0.1⟨*D*_*a*_⟩ = 10 *μm* and standard deviation *δ*_*σ*_/⟨*σ*_*n*_⟩ = 0.1. We set *ε*_*a*_ = 0.05*J* so that it is comparable to the bulk modulus of a lipid droplet (*ε*_*a*_/*a*_*α*,0_∼10M*a*)^57^. Furthermore, we set *ε*_*l*_/*ε*_*a*_ = 0.1, *ε*_*aa*_/*ε*_*a*_ = 0.1, *ε*_*cc*_/*ε*_*a*_ = 0.05, *ε*_*ac*_/*ε*_*a*_ = 0.1, *ε*_*w*_/*ε*_*a*_ = 0.1, *ζ* = 0.01, and *a*_*c*_ = 0.1. We vary *ε*_*b*_/*ε*_*a*_ from 10^-4^ to 10^-2^ to study the effect of adipocyte stiffness on breast cancer invasion.

#### Compressive Stiffness of an Adipocyte

To characterize the compressive stiffness of an adipocyte, we place the adipocyte between two rigid walls. The adipocyte interacts with the walls via *U*_*w*_. We decrease the distance between two walls by a small amount and perform energy minimization for the adipocyte. We then measure the force exerted on the walls. We repeat this process iteratively and determine the compressive stiffness as the slope of the wall force versus the adipocyte compressive strain.

#### Initial Adipocyte-Cancer Cell Packing

For the simulations of cancer cell invasion into adipose tissue, we first generate a static packing of adipocytes and breast cancer cells with an abrupt interface at a target pressure *P*_*t*_. To prevent adipocytes from mixing with cancer cells in the static packing, we initially generate the adipocyte packing and the cancer cell packing separately and then put them together. To generate an adipocyte packing, we start from a dilute packing with packing fraction *φ* = 0.01 in a square box (with the same lengths *L*_*x,a*_ and *L*_*y,a*_ along the x- and y-directions) with two straight walls to confine the packing in the *x*-direction and periodic boundary conditions in the *y*-direction in 2D. We then compress the packing by reducing *L*_*x,a*_ and *L*_*y,a*_ by the same amount with a change in packing fraction Δ*Ø* = 0.001, followed by energy minimization. If *P* is smaller than *P*_*t*_, we compress the packing again. Otherwise, we return to the packing before the compression and compress the system by half of the original Δ*Ø*. We repeat this process until we reach *P*_*t*_ to within *P*_*t*_ ± 0.01*P*_*t*_. We apply the same procedure to generate a packing of breast cancer cells in a box with length in the y-direction *L*_*y,c*_= *L*_*y,a*_, and we only change the box length along the x-direction *L*_*x,c*_ to compress or decompress to achieve *P*_*t*_. We then join the adipocyte and breast cancer cell packings and repeat the compression and decompression procedure by changing only the box length along the x-direction until *P* of the entire packing reaches *P*_*t*_ to within *P*_*t*_ ± 0.01*P*_*t*_.

#### Breast Cancer Invasion

After generating a static packing of adipocytes and cancer cells, we assign velocities to the cancer cells drawn from a Gaussian distribution with zero mean in both the x- and y-directions and a variance determined by the temperature T. We then carry out constant temperature and pressure molecular dynamics simulations. The equations of motion for the cancer cells, adipocytes, and the mobile wall are given by:

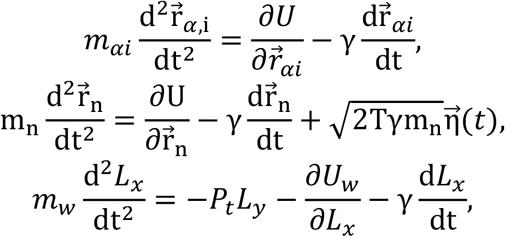

where *m*_*αi*_, *m*_*n*_, and *m*_*w*_ are the masses of vertex i on adipocyte *α*, cancer cell n, and mobile walls, respectively, 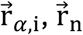, and *L*_*x*_ are the positions of vertex i on adipocyte *α*, cancer cell n, and mobile walls, respectively, γ is the damping coefficient, and 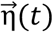 is a two-dimensional Gaussian noise strength that has zero mean and unit standard deviation in both the x- and y-directions and zero covariance. We use the cancer cell diameter (10 *μm*) and mean speed of individual cancer cells in isolation (30 *μm*/*h*) to set the length and time scales of the simulations^58^.

#### Characterization of the Degree of Invasion

We quantify the degree of invasion by calculating the interface length between adipocytes and breast cancer cells. Specifically, we apply the Laguerre-Voronoi tessellation based on the positions of cancer cells and vertices of the adipocytes. After the Laguerre-Voronoi tessellation, a Voronoi polygon is associated with each breast cancer cell and adipocyte vertex. We identify all edges shared between two polygons. We discard edges shared between vertices that belong to the same adipocyte. We define the invasion degree as

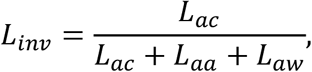

where *L*_*ac*_, *L*_*aa*_, and *L*_*aw*_ are the total length of edges shared between adipocytes and cancer cells, between different adipocytes, and between adipocytes and walls, respectively. We further rescale *L*_*inv*_ by its minimum *L*_*min*_ and maximum *L*_*max*_ so that the scaled invasion length 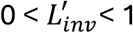:

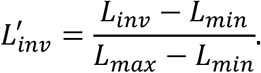

### 3.17 Statistical methods

Statistical significance was evaluated by unpaired two-tailed Student’s *t* test for the comparison of two conditions, and by one-way analysis of variance (ANOVA) followed by Tukey’s test for multiple comparisons. Unless otherwise noted, p < 0.05 for all analyses and significance is depicted by compact letter display notation (i.e., groups with the same letter are not statistically different, while groups with distinct letters are statistically different from each other). Averaged data is reported as the mean plus or minus the standard deviation (SD) unless specified.

## 4. Results

### 4.1 Tunable polyacrylamide beads recapitulate primary adipocyte size and stiffness

Primary adipocytes and a collagen-rich ECM constitute the primary structural components of adipose tissue in the breast^11,12^ (**Fig. 1a, left**). During tumor invasion, breast cancer cells at the invasion front must physically navigate this environment, migrating through the interstitial spaces between individual adipocytes as single cells or in small clusters^5^ (**Fig. 1a, right**). To better understand adipocyte size and deformability as two biophysical features which influence tumor cell migration, we first analyzed histological sections of mammary fat pads collected from female C57BL/6 mice (**Fig. 1a, left**). We observed substantial heterogeneity in the relative size and deformation of densely packed adipocytes, and characterized the cross-sectional areas *A* of individual adipocytes to estimate cell diameters 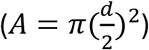. Effective diameters ranged from d = 6.70μm to 65.2μm, with a mean ⟨*d*⟩ = 31.2μm and standard deviation Δ*d* = 9.78μm (**Fig. 1b**). To confirm that these two-dimensional size measurements could be used to approximate three-dimensional trends, we show that ⟨*d*⟩ is monotonically related to the true average diameter of the adipocytes, ⟨*D*⟩, and the standard deviation *σ*_*d*_ of d is proportional to the standard deviation *σ*_*D*_ of true diameter *D* (**Fig. S1a**). In addition, we isolated primary adipocytes from a subset of resected tissue^43^ for mechanical characterization via atomic force microscopy (AFM), immobilizing the cells on the imaging substrate during data acquisition (setup shown in **Fig. S1b**). Mirroring the qualitative observation of variable adipocyte deformation in tissue sections, we found primary adipocyte stiffness to be considerably heterogeneous, with the elastic moduli of cells ranging from a few hundred to several thousand pascals (**Fig. 1c**). This broad range of values additionally aligns with the mechanical variability previously observed with other adipose model systems, though our measurements ranged from similar (100-1000 Pa) to an order of magnitude greater than (∼10,000 Pa) previous reports which used differentiated preadipocyte cell lines^14,15,59^ or long-term explanted adipocyte cultures^60^. These include a study which used the specific AFM protocol and parameters employed in this report^15^, suggesting that our findings are not an artifact of the method but a true feature of the tissue. Collectively, our results reflect an inherent variability of adipocyte stiffness, warranting further investigation into how this heterogeneity influences breast cancer invasion.

**Figure 1:**
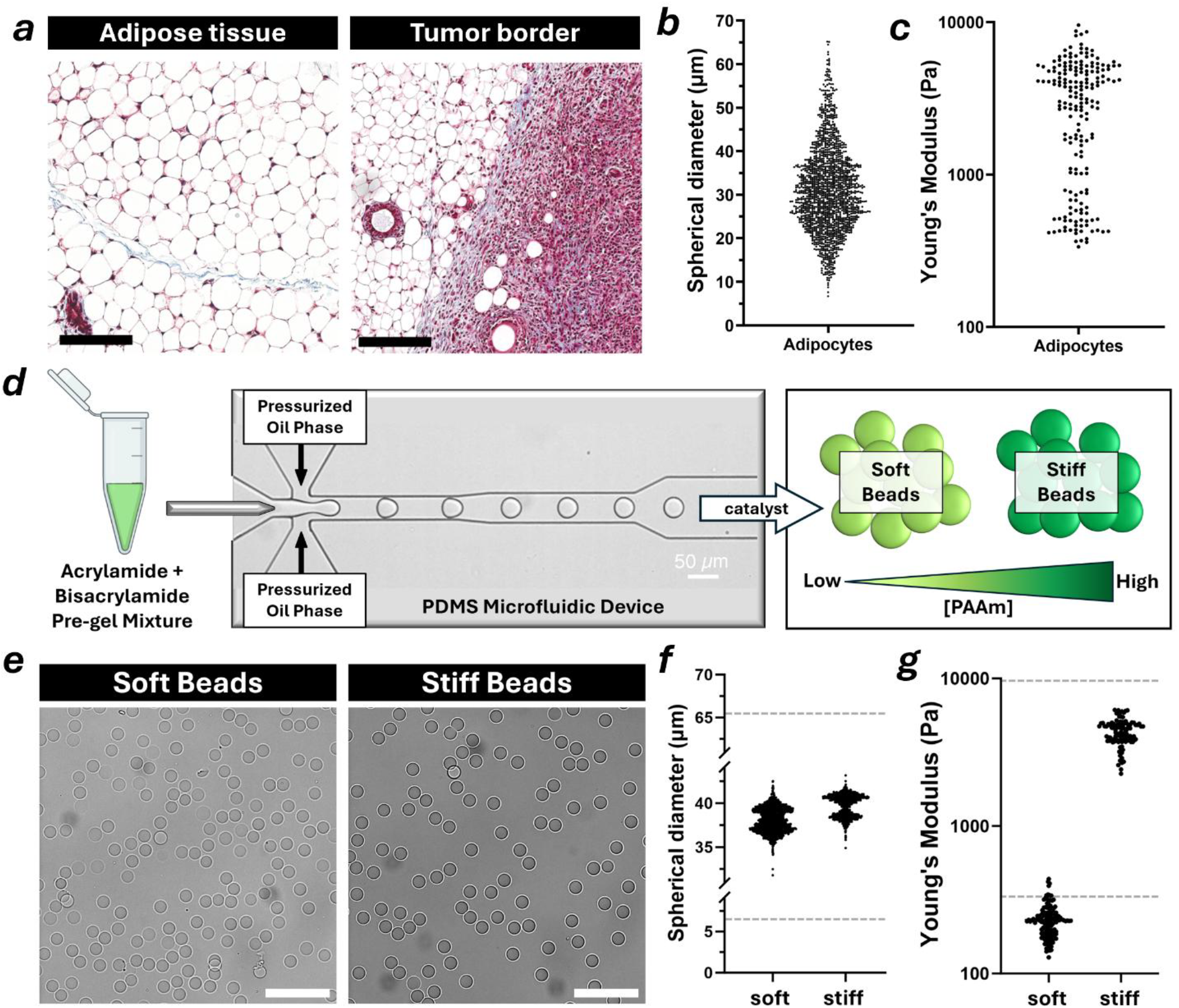
Tunable polyacrylamide beads recapitulate primary adipocyte size and stiffness. **(*a*)** Masson’s trichrome staining of murine adipose tissue, scale = 100 μm. **(*b*)** Murine adipocyte diameter distribution. **(*c*)** Atomic force microscopy analysis of murine adipocyte stiffness distribution. **(*d*)** Schematic of tunable polyacrylamide bead fabrication via custom droplet microfluidics. **(*e*)** Brightfield images of soft and stiff polyacrylamide bead batches, scale = 200 μm. **(*f*)** Polyacrylamide bead diameter distributions, dashed lines represent adipocyte range. **(*g*)** Atomic force microscopy analysis of polyacrylamide bead stiffnesses, dashed lines represent adipocyte range.

To develop a micromechanical, adipose tissue-mimetic model informed by our analysis of adipocyte size and stiffness, we next replicated the physical properties of adipocytes with biochemically inert PAAm beads. We generated these beads using a customizable flow-focusing droplet microfluidic device and restricted the resulting microgel size to approximately 35 μm in diameter, mimicking primary adipocytes, via selective control of the oil phase flow rate. Moreover, bead stiffnesses of about 200 Pa (soft adipocytes) and 4000 Pa (stiff adipocytes) were achieved by altering the concentration of fixed-ratio acrylamide monomers prior to polymerization within a catalyst-containing oil phase^40,41^ (**Fig. 1d**). Using this platform, we were able to generate bead populations of variable stiffness while maintaining nearly monodisperse size distribution (**Fig. 1e**), achieving average diameters comparable to primary adipocytes (**Fig. 1f**). Moreover, AFM characterization of the two bead populations confirmed that their elastic moduli were reflective of the lower (soft beads, ⟨*Y*⟩ = 230 Pa on average) and upper (stiff beads, ⟨*Y*⟩ = 4339 Pa on average) ends of the stiffness values measured for primary adipocytes (**Fig. 1g**). Collectively, these results demonstrate our ability to generate distinct adipocyte-sized populations of PAAm beads with elastic moduli spanning the stiffness heterogeneity of primary adipocytes.

### 4.2 PAAm bead granular hydrogels model the structural organization of adipose tissue

After validating that polyacrylamide microgels can mimic physical properties of primary adipocytes, we next characterized the ECM structure of native adipose tissue. Second harmonic generation (SHG) microscopy of human adipose tissue showed organized collagen fibers occupy the interstitial spaces between individual adipocytes in human adipose tissue (**Fig. 2a**), an arrangement necessary for tumor cell invasion^20,61^. To recreate this microarchitecture, we encapsulated soft or stiff PAAm beads in either a 2.5 mg/mL or 6.0 mg/mL type I collagen hydrogel, reflective of healthy or fibrotic breast tissue phenotypes respectively^9,62,63^. To this end, bead-laden collagen hydrogels, referred to as ‘granular hydrogels’, were cold cast into custom PDMS microwells, as previously described^44^, to encourage collagen fibrillogenesis representative of native ECM microarchitecture (**Fig. 2b**). To develop these granular hydrogels, we trialed collagen-to-bead volume ratios of 1:3, 1:6, and 1:9, and used a 1:6 ratio for all future experiments as this yielded the packing fraction closest to native adipose tissue^11^ while maintaining sufficient collagen content to ensure the structural integrity of the hydrogel (**Fig. S1c-d**). Confocal imaging revealed that interconnected networks of collagen fibers formed between fluorescently labeled PAAm beads yielding architectures reminiscent of interstitial adipose ECM *in vivo* (**Fig. 2c**). While the local bead packing fraction varied within and between individual gels, no global differences were detected across the four combinations of bead stiffness and collagen concentration tested (**Fig. 2d-e**). Hence, any differences detected in subsequent experiments can be attributed to the inherent mechanical properties of the hydrogel rather than differences in porosity. Finally, we confirmed that incorporation of PAAm beads did not impact the concentration of the interstitial collagen phase compared to respective beadless hydrogels (**Fig. 2f**) using an adapted version of a commercially available bicinchoninic acid (BCA) assay (**Fig. S1e-g**). These results ensured that any differences in data collected from the granular hydrogels are not due to variation in collagen density between conditions. Taken together, these findings validate that our model recapitulates key structural characteristics of adipose tissue, and enables control over the packing of adipocyte-mimetics within a selectively tunable ECM.

**Figure 2:**
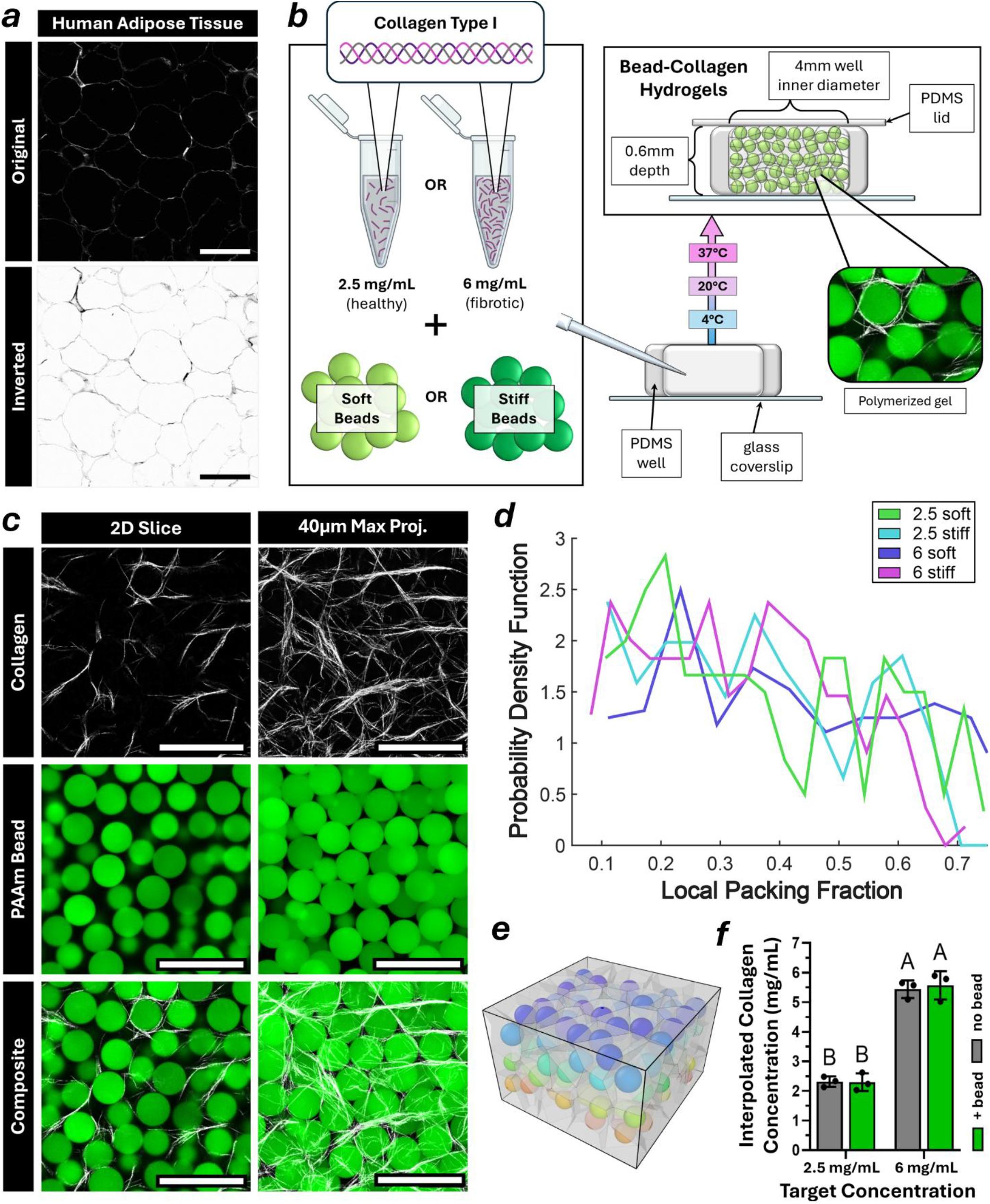
PAAm bead granular hydrogels model the structural organization of adipose tissue. **(*a*)** Second harmonic generation images of interstitial ECM in whole human adipose tissue, scale = 100 μm. **(*b*)** Collagen-embedded polyacrylamide bead hydrogel fabrication schematic. **(*c*)** Reflectance (top) and fluorescence (middle) confocal microscopy images of soft polyacrylamide beads embedded within 2.5 mg/mL collagen, scale = 100 μm. **(*d*)** Analysis of the local packing fraction of polyacrylamide beads in hydrogels of different component stiffnesses. **(*e*)** 3D reconstruction of granular hydrogel bead packing. **(*f*)** Bicinchoninic acid assay analysis of type I collagen concentration in beadless and bead-embedded hydrogels.

### 4.3 PAAm beads promote cancer cell invasion through collagen in a stiffness-dependent manner

Next, we assessed how altering bead stiffness and collagen concentration impacts breast cancer cell migration, to approximate the mechanical regulation of breast cancer cell invasion by native adipose tissue. To this end, we seeded invasive MDA-MB-231 triple-negative breast cancer cells on the exposed surface of granular hydrogels and assessed their invasion into the hydrogels after 36 hours via confocal microscopy (**Fig. 3a**). This experimental configuration mimicked the interactions of single or small clusters of cancer cells with the native stromal interface at the leading edge of a tumor, independent of tumor growth-induced mechanical force such as solid stress or interstitial fluid pressure (IFP)^64^. Cells invaded into the hydrogel in both soft and stiff bead conditions (**Fig. 3b**) and appeared to organize and elongate around PAAm beads. While no significant trends in cell morphology were observed between conditions, the inclusion of either soft or stiff beads significantly increased the depth of tumor cell invasion compared to beadless hydrogels (**Fig. 3c**). These findings were highly consistent between experimental replicates (**Fig. S2a**). Additionally, nuclear tracking revealed that MDA-MB-231s invaded the most in 2.5 mg/mL gels with soft beads and the least in 6.0 mg/mL beadless controls, with soft beads facilitating migration compared to stiff or no beads and 2.5 mg/mL collagen facilitating migration compared to 6.0 mg/mL collagen (**Fig. 3d-e**). Interestingly, the mean invasion depths of the two intermediate granular hydrogels containing both soft and stiff tissue components (2.5 mg/mL collagen + stiff beads and 6 mg/mL collagen + soft beads) were similar, but the distribution of cell invasion differed between both systems, suggesting concurrent but independent regulation of cell invasion by collagen and beads. These findings were not an artifact of the cell line chosen as we observed similar trends in a subset of conditions with human MCF10A mammary epithelial cells and murine EO771 hormone receptor-positive breast cancer cells, supporting the broader relevance of our findings (**Fig. S2b-c**). Specifically, bead inclusion within the collagen ECM promoted invasion in a stiffness-dependent manner across all cell lines regardless of baseline motility, demonstrating that adipocyte-mimetic bead stiffness and ECM density broadly dictate the migratory activities of mammary cells. Finally, to better understand the migratory behavior of tumor cells in granular hydrogels, we estimated tumor cell diffusion constants for each condition by plotting the corresponding invasion depth distribution against a half-normal probability distribution, which is known to model diffusive cell motion^55^. Diffusion constants mirrored absolute invasion depth measurements (**Fig. 3f**), though the migratory behavior of cells in stiff bead or beadless conditions was less accurately captured by a diffusive model than cells in soft bead conditions, as indicated by diverging R^2^ values (**Fig. S2d-e**). We propose these results stem from additional physical constraints imposed on cells by stiffer beads or denser collagen which restrict tumor cell migration and limit diffusive motion. Collectively, our results indicate that the inclusion of PAAm beads facilitates breast cancer cell migration into collagen hydrogels, with the most pronounced effects occurring in low density collagen hydrogels containing soft beads.

**Figure 3:**
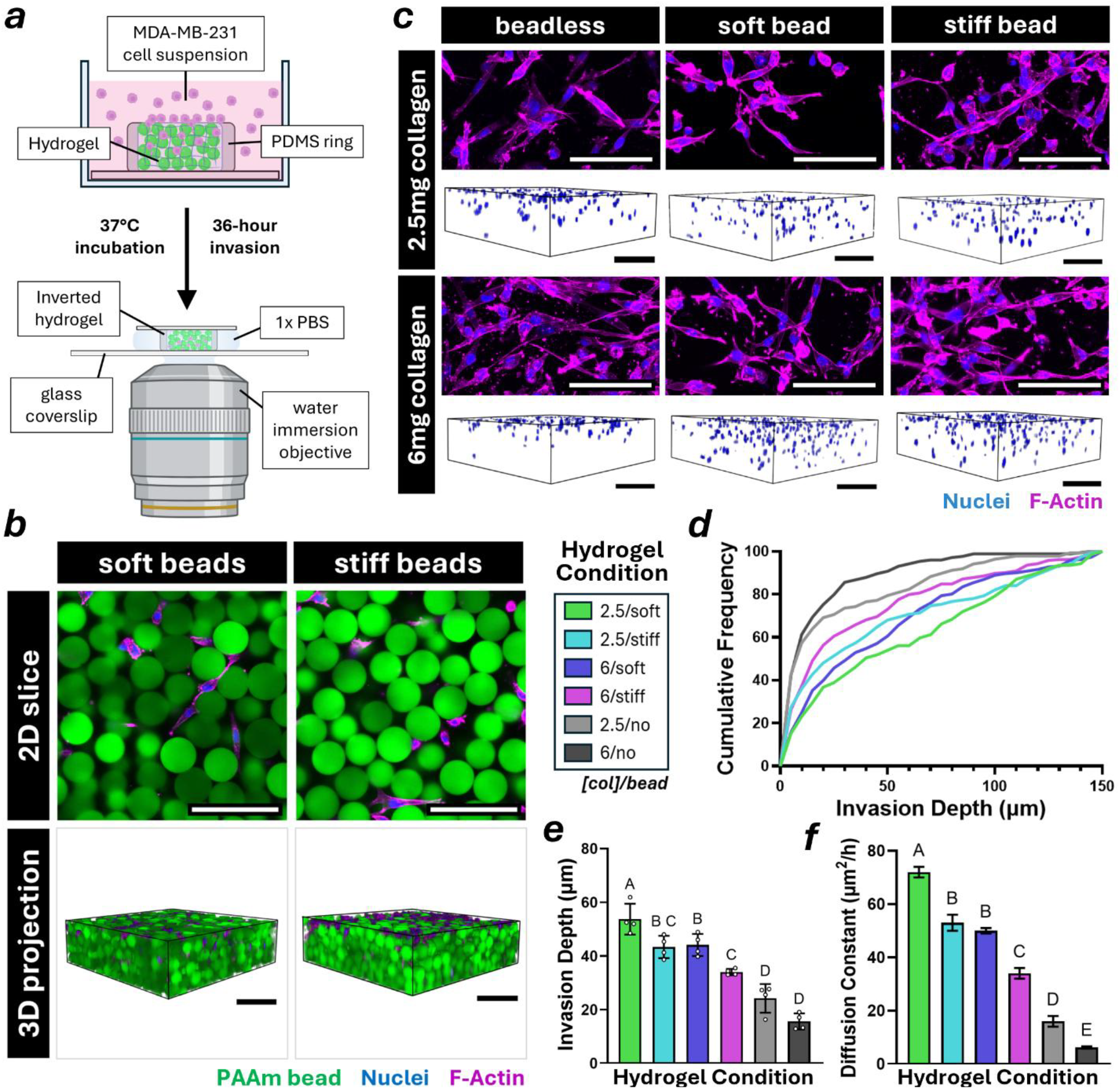
PAAm beads promote cancer cell invasion through collagen in a stiffness-dependent manner. **(*a*)** Schematic of MDA-MB-231 cancer cell invasion assay experimental setup. **(*b*)** Confocal micrographs of bead-laden 6 mg/mL collagen hydrogels seeded with MDA-MB-231 breast cancer cells stained with DAPI (blue) and phalloidin (magenta), scale = 100 μm. **(*c*)** Confocal micrograph 150 μm maximum projections (top) and 3D projections (bottom) of MDA-MB-231 cells invading through 2.5 or 6 mg/mL beadless and granular collagen hydrogels, scale = 100 μm. **(*d*)** Cumulative frequency distribution of MDA-MB-231 cell invasion depth after 36-hour invasion through granular hydrogels of differing stiffness. **(*e*)** Mean ± standard deviation MDA-MB-231 cell invasion depth after 36-hour invasion through granular hydrogels of differing stiffness. **(*f*)** Mean ± standard deviation MDA-MB-231 cell diffusion constants.

### 4.4 Inclusion of PAAm beads alters collagen fiber architecture in a concentration-dependent manner

We next sought to characterize how bead incorporation affected collagen network formation, considering its established role in regulating tumor cell migration^65,66^. In native adipose tissue, collagen fibers exhibit linear and aligned geometry between adipocytes^19^, Moreover, it has been shown for other biopolymers that milieu-dependent changes in assembly kinetics influence filament elongation and lateral association^67^. Therefore, we hypothesized that bead incorporation alters collagen fibrillogenesis and anisotropy compared to bead-free collagen. Indeed, confocal reflectance microscopy revealed that inclusion of beads promoted the formation of hierarchical networks of aligned collagen fibers, while beadless hydrogels of both collagen densities lacked this organization (**Fig. 4a)**. Collagen fibers were also slightly thinner in beadless conditions and there were no appreciable differences in fiber length or fiber straightness (**Fig. S3a**). To quantify potential differences in fiber alignment more rigorously, we used the open-source Fiji plugin OrientationJ^49^ which can assign each collagen fiber in an image a coherency score based on its relative alignment to nearby fibers (**Fig. 4b**). Notably, collagen fibers in hydrogels with PAAm beads were significantly more coherent, or anisotropic at a locally defined scale, than fibers in beadless hydrogels (**Fig. 4c**) with high consistency between experimental replicates (**Fig. S3b**). Of note, 2.5 mg/mL hydrogels demonstrated a higher fraction of aligned collagen fibers compared to 6.0 mg/mL hydrogels, regardless of whether PAAm beads were included (**Fig. 4d**). Although bead stiffness did not affect fiber coherency between collagen density-matched granular hydrogels, we observed a trend suggesting an inverse relationship between bead stiffness and average fiber alignment (**Fig. S3c**). As further proof of concept, we quantified the eccentricity of individual collagen fibers computationally^48^, and observed similar trends whereby fiber eccentricity increased upon bead integration compared to beadless controls (**Fig. S3d-e**). Moreover, considering the preferential migration of cells along highly aligned ECM structures^65^, we aimed to assess the three-dimensional interconnectivity of collagen fibers as a function of fiber alignment. To this end, we employed a custom image analysis workflow which allowed us to isolate grouped collagen fibers in accordance with an increasingly stringent coherency threshold (**Fig. S3f**). This analysis yielded volumetric and raw count data for both the complete collagen network as well as smaller, interconnected collagen networks identified above a certain coherency threshold. These data were then used to define a ‘connectivity’ term describing the thresholded collagen network volume as a fraction of the total unthresholded collagen network volume (**Fig. 4e**). We found that the collagen fibers in granular hydrogels were significantly more interconnected than those of beadless gels across all coherency scores which surpassed the percolation threshold (**Fig. 4e**). At higher coherency thresholds, granular hydrogels similarly demonstrated increased fiber interconnectivity when fabricated with 2.5 mg/mL collagen compared to 6.0 mg/mL collagen, suggesting that the hierarchical structure of collagen networks is coordinated by bead inclusion and is dependent on local ECM density. Collectively, our results indicate that the inclusion of PAAm beads coordinates collagen fiber architecture in granular hydrogels, yielding more aligned and interconnected fibers relative to beadless controls, an effect that is more pronounced effects at lower collagen densities.

**Figure 4:**
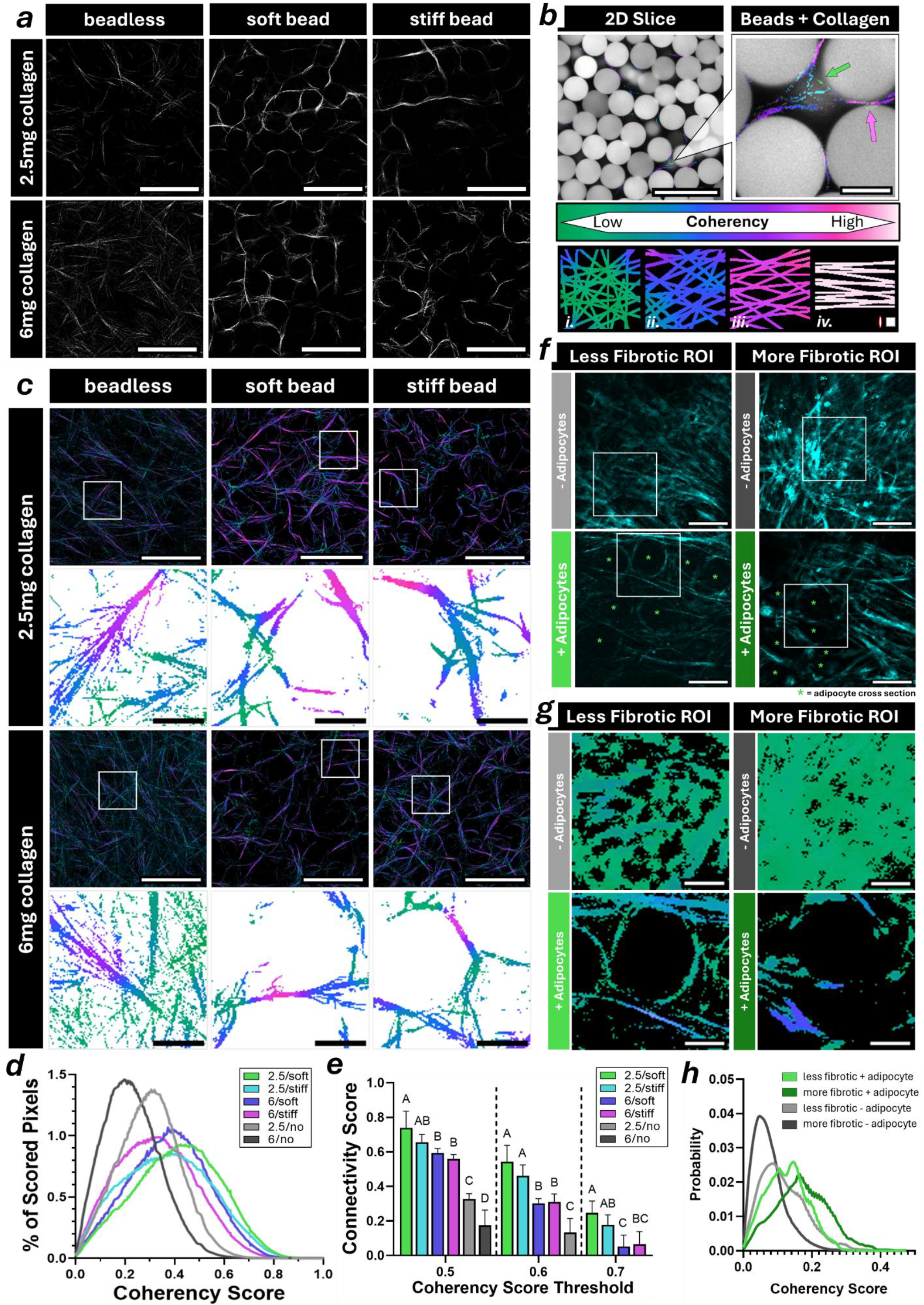
Inclusion of PAAm beads alters collagen fiber architecture in a concentration-dependent manner. **(*a*)** 2D slice reflectance micrographs of 2.5 or 6 mg/mL type I collagen in beadless and granular hydrogels, scale = 100 μm. **(*b*)** Collagen fiber coherency color survey analysis, and representative networks of variable alignment. Scale = 100 μm (left), 25 μm (right), focal adhesion scale approximation = 1x4 μm (red/white ellipse, bottom right), window size = 3 μm (white square, bottom right). **(*c*)** 40 μm maximum projection reflectance micrographs of 2.5 and 6 mg/mL type I collagen overlaid with coherency score color survey (top) and 2D slice raw color survey output (bottom), scale = 100 μm (top), 20 μm (bottom). **(*d*)** Condition-dependent fiber coherency score distributions for granular hydrogels. **(*e*)** Condition-dependent fiber connectivity analysis, grouped by coherency score threshold. Statistics calculated within threshold groups only. **(*f*)** Intravital 2-photon microscopy of collagen shown by second harmonic generation (SHG), compiled into 10 μm maximum projections capturing varying levels of fibrosis and adipocyte packing in live murine mammary tissue, scale = 25 μm, square ROI zoom corresponding to **(*g*)**, Coherency color survey analysis of 10 μm maximum projection SHG micrographs depicting collagen fiber structure in living murine mammary fat pads imaged via intravital microscopy, zoomed scale = 10 μm. **(*h*)** Condition-dependent fiber coherency score distributions for living murine mammary fat pads.

**Figure 5:**
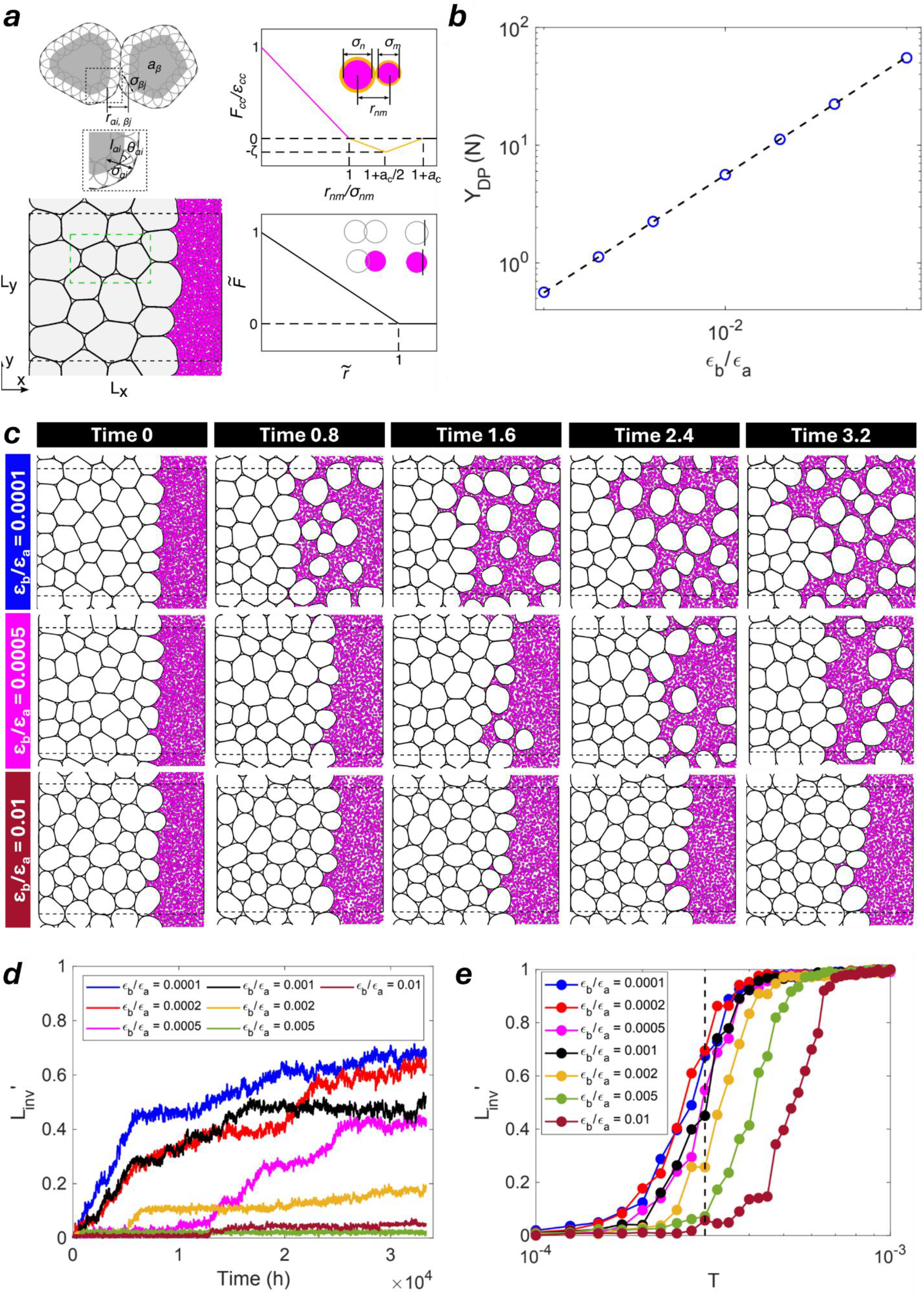
Discrete Element Method (DEM) modeling suggests adipocyte stiffness regulates breast cancer progression. **(*a*)** DEM simulations of breast cancer cell (magenta disks) invasion into packings of adipocytes (grey deformable polygons) in two dimensions. Adipocytes are modeled as deformable particles (top left panel). Cancer cells are modeled as soft disks with an adhesive shell (top right panel, where inter-cancer cell forces *F*_*cc*_ are calculated as the negative gradient of the potential energy *U*_*cc*_). The packings of cancer cells and adipocytes are confined to a rectangular box with periodic boundary conditions in the y-direction and two confining walls at x=0 and x=L_x_ (bottom right panel). A schematic of the purely repulsive force for adipocyte-adipocyte 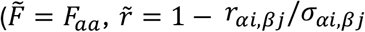, overlapping gray disks), adipocyte-cancer 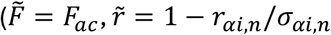 overlap between gray and magenta disks), adipocyte-wall 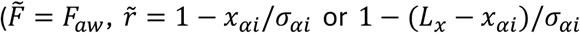, overlap between gray disk and vertical line), and cancer-wall 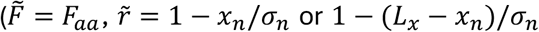, overlap between magenta disk and vertical line) interactions is provided in the bottom right panel. **(*b*)** The compressive stiffness of a single adipocyte Y_DP_, versus the bending strength ε_*b*_, normalized by ε_*a*_, using the deformable particle model. **(*c*)** Examples of p_4_ ackings of cancer cells and adipocytes with different bending energies ε_*b*_/ε_*a*_ at different times (x 10 hours) during invasion. **(*d*)** Rescaled invasion degree 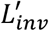 plotted versus time for various ε_*b*_/ε_*a*_ indicated by the colors. **(*e*)** The invasion degree 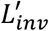 at long times plotted versus the cancer cell temperature T for various ε_*b*_/ε_*a*_ indicated by the colors. Three different configurations are used for average for each ε_*b*_/ε_*a*_.

To validate the *in vivo* relevance of these findings, we employed high resolution multiphoton imaging techniques on the mammary fat pads of live mice^45,46^. Second Harmonic Generation (SHG, collagen) images depicting tissue regions varying in levels of collagen fibrosis and local adipocyte packing were acquired (**Fig. 4f, S4a**). Comparing the ratios of positive (collagen signal) to negative (background) pixel values between conditions demonstrated differing degrees of fibrosis represented by the image stacks selected for analysis (**Fig. S4b**). Next, to compare relative levels of fiber alignment dependent on adipocyte presence and fibrosis, these image stacks were subjected to the same OrientationJ color survey workflow as the *in vitro* image sets (**Fig. 4g, S4c**). Mirroring the trends demonstrated by the granular hydrogels, adipocyte packing appeared to coordinate fiber architecture compared to paired tissue regions absent of adipocytes; fiber alignment increased around adipocytes regardless of fibrosity, corresponding to higher bulk tissue coherency scores (**Fig. 4h**). We note that raw coherency scores differ greatly between *in vitro* and *in vivo* data, and attribute this variability to differences in image acquisition parameters such as objective magnification and step size, characteristically poor intravital resolution due to light scattering, and the presence of other tissue structures such as vasculature. Interestingly, however, the coherency scores of fibrotic conditions appeared to increase more drastically in the presence of adipocytes compared to less fibrotic tissue regions, again paralleling the trends demonstrated in the granular hydrogels (**Fig. 4h**).

### 4.5 Discrete Element Method (DEM) modeling suggests adipocyte stiffness regulates breast cancer progression

We observed that breast cancer cell invasion increased in soft bead hydrogels compared to stiff bead hydrogels, regardless of collagen concentration, and that collagen fiber alignment did not significantly differ with bead stiffness. Given that differences in collagen architecture thus cannot explain the observed changes in invasion, we wanted to explore the effect of bead stiffness. As such, we employed Discrete Element Method (DEM) simulations to computationally demonstrate how individual adipocyte stiffness may inform bulk breast cancer progression. First, we varied the stiffness of the adipocytes via the bending strength ε_*b*_ to study the impact of adipocyte stiffness on breast cancer invasion. We found that the compressive stiffness of a single adipocyte increases linearly with ε_*b*_ (**Fig. 5b**). Next, in typical simulations of breast cancer invasion into adipose tissue, we found that the degree of invasion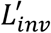 increases over time and then reaches a plateau at long time points, as shown in **Fig. 5c-d**. The long-time plateau in 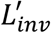 increases with increasing mean speed or activity as parameterized by T (**Fig. 5d**), in agreement with experimental observations that more motile cancer cells are more invasive. Furthermore, at the same of activity T, we found that cancer cells are less invasive when navigating stiffer adipocytes (**Fig. 5d**) and that cancer cells must be more motile to achieve the same degree of invasion 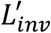 into adipose tissue with stiffer adipocytes compared to that for softer adipocytes (**Fig. 5e**). The observation of hindered cancer invasion into adipose tissue with increased adipocyte stiffness is consistent with the experimental results presented above. We note that the predicted timescale for full invasion of the cancer cells into adipose tissue is much longer compared to that found in experiments (less than 2 days). This difference can be attributed to the fact that we use the typical invasion speed of cancer cells in isolation in this model^58^, and future intravital time course studies should be conducted to better characterize the velocity of cells *in vivo*.

### 4.6 PAAm bead-induced cell nuclear deformation and local packing geometry influence breast cancer cell invasion

Physical constraints regulate tumor cell invasion due, in part, to the rate-limiting process of nuclear deformation during confined migration^68,69^. To assess how the organization and confined packing of PAAm beads impacted migration dynamics in our system, we tracked the trajectories of invading MDA-MB-231 breast cancer cells through the most mechanically permissive granular hydrogel condition (2.5 mg/mL collagen + soft beads) (**Fig. 6a**). While cancer cell migration speeds fluctuated significantly over the invasion period, no generalizable correlation between cell speed and bead proximity was discernible. However, when tracking only the most motile cells, we observed that these subpopulations stalled, rapidly accelerated, and then decelerated as they squeezed between adjacent beads (**Fig. 6b, S5a**). This pattern aligns with previous studies of confined migration, which showed that tumor cells produce cytoskeletal protrusions to pull themselves through confining spaces^70,71^, consequentially deforming their nuclei and generating additional forces through a recoil mechanism, before returning to slower migration speeds^72^. In line with this model, we observed both bead (**Fig. 6c**) and nuclear deformation (**Fig. 6d**) when migrating tumor cells were in direct contact with the adipocyte-mimetic PAAm beads. Specifically, both soft and stiff beads were significantly deformed at the cell contact interface compared to those uncontacted by cells (**Fig. 6e**). Similarly, nuclei localized in unconfined void spaces between beads were significantly rounder than nuclei between bead-bead interfaces, while nuclei in beadless hydrogels adopted an intermediate phenotype (**Fig. 6f**). Moreover, tumor cells were able to deform soft beads significantly more than stiff beads, consistent with a less physically restrictive environment and the greater cell invasion depths we observed in soft granular hydrogels. Interestingly, we did not detect differences in bead deformation between 2.5 mg/mL and 6.0 mg/mL granular hydrogels (**Fig. S5b**), highlighting the need to consider the mechanics of other tissue features besides the ECM. Likewise, we did not observe consistent differences in nuclear deformation across mechanical conditions (**Fig. S5c**), indicating that nuclear stiffness may be a limiting factor in the migration trajectories we measured. Taken together, these data suggest that both tumor cell intrinsic (e.g. nuclear^68,69,73^ or cell stiffness^74,75^) and extrinsic (bead/adipocyte) mechanics influence the migratory behavior of breast cancer cells in a dynamic, multi-step manner (**Fig.6g**).

**Figure 6:**
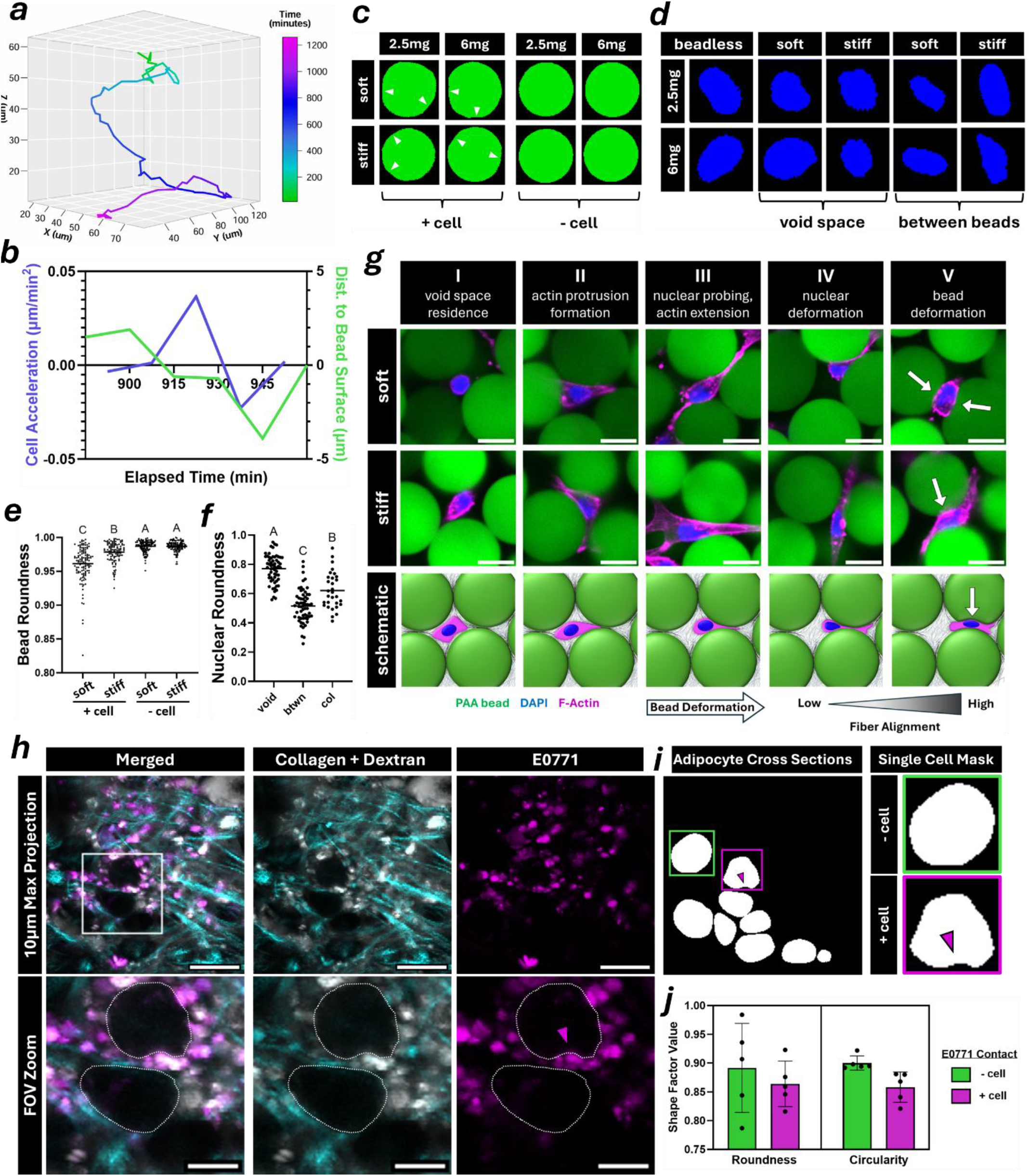
PAAm bead-induced cell nuclear deformation and local packing geometry influence breast cancer cell invasion. **(*a*)** Plot of a single tracked nucleus migrating through a soft bead 2.5 mg/mL granular hydrogel imaged over 21 hours via spinning disk confocal microscopy. **(*b*)** Plot depicting a moment of nuclear acceleration (left axis, blue) correlating with a reduction in nearest bead proximity (right axis, green) as a single cell squeezes between two beads. **(*c*)** Segmented masks of 2.5 or 6 mg/mL granular hydrogel bead deformation with or without cell contact. **(*d*)** Segmented cell masks of nuclear deformation for cells residing in the void or interstitial spaces between beads in 2.5 or 6 mg/mL granular hydrogels. **(*e*)** Plot of soft and stiff PAAm bead deformation, collected from both 2.5 and 6 mg/mL granular hydrogels, evaluated via roundness shape parameter. **(*f*)** Plot of nuclear deformation data, collected from both 2.5 and 6 mg/mL collagen hydrogels, describing the localization of a nucleus to unconfined void regions (‘void’) or confined regions between (‘btwn’) soft or stiff PAAm beads in granular hydrogels, or in beadless collagen (‘col’) hydrogels, evaluated via roundness shape parameter. **(*g*)** Schematic depicting migratory mechanisms of single cells invading between adipocyte-mimetic beads with corresponding micrographs captured in 2.5 mg/mL collagen granular hydrogels, scale = 25 μm. **(*h*)** Intravital micrographs compressed into 10 μm maximum projections of SHG (collagen) and fluorescent (separate dextran and E0771) signal, depicting breast cancer cells (magenta) migrating between and deforming adipocytes (gray outline) in living murine mammary tissue, scale = 25 μm (top), 10 μm (bottom), arrows indicate E0771 contact deformation. **(*i*)** Manually interpolated single slice cell masks of live mammary tissue-resident adipocytes (left), with representative single-cell masks depicting cancer cell-deformed (pink/bottom) and undeformed (green/top) adipocyte cross sections, arrows indicate E0771 contact deformation. **(*j*)** Graph depicting differences in adipocyte cross section roundness and circularity depending on E0771 cell contact.

To determine how invasive tumor cells affect adipocyte morphology *in vivo*, intravital images of murine mammary fat pads were acquired two weeks after being orthotopically injected with syngeneic EO771-YFP cancer cells. By co-registering images of cancer cells with high-contrast slices of surrounding collagen (SHG) and blood plasma (fluorescent dextran), we manually identified adipocytes that did or did not contribute to the confinement of invading EO771 cells (**Fig. 6h**, see Materials & Methods for selection criteria). Consistent with our *in vitro* observations, cancer cell confinement between adipocytes correlated with adipocyte deformation (**Fig. 6i**). Quantification of adipocyte shape via image analysis confirmed that adipocytes in direct contract with EO771 cells were less circular and round as compared to adipocytes not in contact tumor cells (**Fig. 6j**). Adipocytes in contact with relatively unconfined cancer cells also exhibited some level of deformation, but not to a comparable degree (**Fig. 6i**). The parallels between these *in vivo* findings and our *in vitro* data collected from the granular hydrogels suggest that mechanical interactions between cancer cells and adipocytes may play a role in tumor dissemination through the adipocyte-rich mammary stroma.

## Discussion & Conclusions

We developed mechanically tunable granular hydrogels to mimic structural features of adipose tissue and demonstrated their utility in studying the physical regulation of breast cancer invasion. More specifically, we fabricated mechanically distinct populations of polyacrylamide beads^40,41^ to use as micromechanical models of primary adipocytes. We incorporated these beads into 3D collagen hydrogels to replicate the physical constraints that breast cancer cells experience as they invade between mammary adipocytes and their surrounding ECM^5,11,12^. The presence of beads increased 3D cell invasion in a manner that depended on bead stiffness and collagen density. Given the established role of fibrillar ECM assembly dynamics^76^ and local architecture^20,65,66^ in the regulation of cell migration, we analyzed the fibrous collagen networks and found that PAAm bead integration significantly altered local scaffold architecture. Finally, considering that physical confinement regulates cancer cell malignancy^77,78^ and migration speed^21,70,72^, we employed a DEM model of breast cancer-seeded adipose tissue to demonstrate that individual adipocyte stiffness dictates the degree of cancer cell invasion through that tissue. We validated these findings both *in vitro* using tissue-mimetic granular hydrogels of varying bead stiffness, and *in vivo* via intravital mammary fat pad imaging revealing that reciprocal deformation of migratory cell nuclei and PAAm beads dictated invasion speed in a stiffness-dependent manner. Taken together, our data collected from these experiments suggest that adipose tissue structure and mechanical properties regulate breast cancer invasion through mammary stroma.

First and foremost, we observed a significant increase in breast cancer cell invasion through bead-laden granular hydrogels compared to beadless controls and similarly found that both soft beads and dilute ECM promoted invasion compared to their stiffer counterparts. Interestingly, cells invading through soft bead granular hydrogels demonstrated a more diffusive phenotype compared to stiff bead granular hydrogels and beadless controls, both of which promoted cell invasion patterns that could not be attributed to diffusion alone. These data suggest that adipocyte structure physically coordinates the tumor microenvironment in a manner which promotes interstitial cancer cell migration. Stiffer tissue components then force invasive tumor cells to mechanically adapt to their local microenvironment, mirroring the findings of several previous studies which demonstrated that tissue organization^79^ and decreased 3D substrate stiffness^80,81^ promote cell migration and select for aggressive metastatic phenotypes^82^.

In tandem, our analyses highlighted an ECM density-dependent increase in local fiber alignment and hierarchical network structure in granular hydrogels compared to beadless controls. Considering the well-characterized role of local ECM anisotropy in the directional contact guidance of migratory cells^20,65,66^, these data suggested that our cell invasion findings could be attributed, at least in part, to bead-coordinated differences in fiber structure. In support of this theory, intravital imaging of EO771 breast cancer-seeded murine mammary fat pads revealed similar patterns occurring *in vivo*; tissue regions with ranging degrees of fibrosis demonstrated increases in collagen fiber organization when packed with adipocytes. Finally, we observed no mechanically coordinated differences in the nuclear shape factor of migrating cells, indicating that adipocyte mechanics do not inherently alter nuclear deformability. PAAm beads in direct contact with cells, however, demonstrated significant stiffness-dependent differences in deformation, which corresponded with moments of nuclear acceleration and deceleration as migratory cells squeezed between beads. Cell nuclei, which are traditionally considered to be the stiffest organelle^83^, are known to act as a rate-limiting factor in confined cell migration in order to prevent DNA damage^68,69,84^. Within this context, our data suggests that as breast cancer cells migrate between adipocytes during invasion, adipocyte stiffness-coordinated confinement of migratory nuclei dictates the velocities of the tumor cell. This data also mirrors the findings of other studies which suggest that some degree of nuclear confinement temporarily alters cell migration through a force-generating recoil mechanism^72^. We therefore posit that the mechanical properties of stiff adipose tissue, while restrictive of diffusive cell movement, may further coordinate the migration of invasive cancer cells by promoting such moments of nuclear recoil.

While the intravital data presented in this report did not have the resolution to evaluate such patterns in invasive cell nuclei, we were able to visualize the reciprocal deformation demonstrated by confinement-inducing adipocytes. Thus, to validate these reciprocal cell-bead deformation patterns *in vivo*, the shape factors of individual adipocyte cross sections that were or were not contributing to the confinement of cancer cells were compared. This analysis yielded similar results to our *in vitro* experiments, demonstrating increased deformation of adipocytes contributing to cancer cell confinement, and further suggesting that adipocyte packing in mammary tissue physically regulates the invasive activity of breast cancer. Additional studies will be needed to explore this hypothesis and elucidate whether additional tumor cell mechanical characteristics influence adipose tissue invasion, especially considering the recent demonstration of how breast tumor cell viscosity dictates interactions with the vasculature in downstream metastatic processes^85^.

Given our findings using PAAm beads as adipocyte mimetics, we propose the following theoretical model of breast cancer cell invasion through adipose tissue (**Fig. 6g, 7a-c**). First, a given tumor cell resides within a less confined void space of the tissue (I), followed by the formation of actin-rich protrusions which extend along the dominant axis of collagen fiber alignment between adjacent adipocytes (II). These protrusions extend beyond this region of peak confinement, then begin to pull the tumor cell through the interstitial space between adipocytes, with the nucleus acting as a mechanical probe limiting migration (III). The migrating nucleus begins to deform, generating recoil force until it achieves a certain threshold at which it can no longer be compressed without risk of DNA damage due to confinement-induced chromatin remodeling (IV), which could ultimately lead to adaptive differences in tumor cell aggression. Finally, the tumor cell rapidly passes between the adipocytes which reciprocally deform in response to transmigration (V).

**Figure 7:**
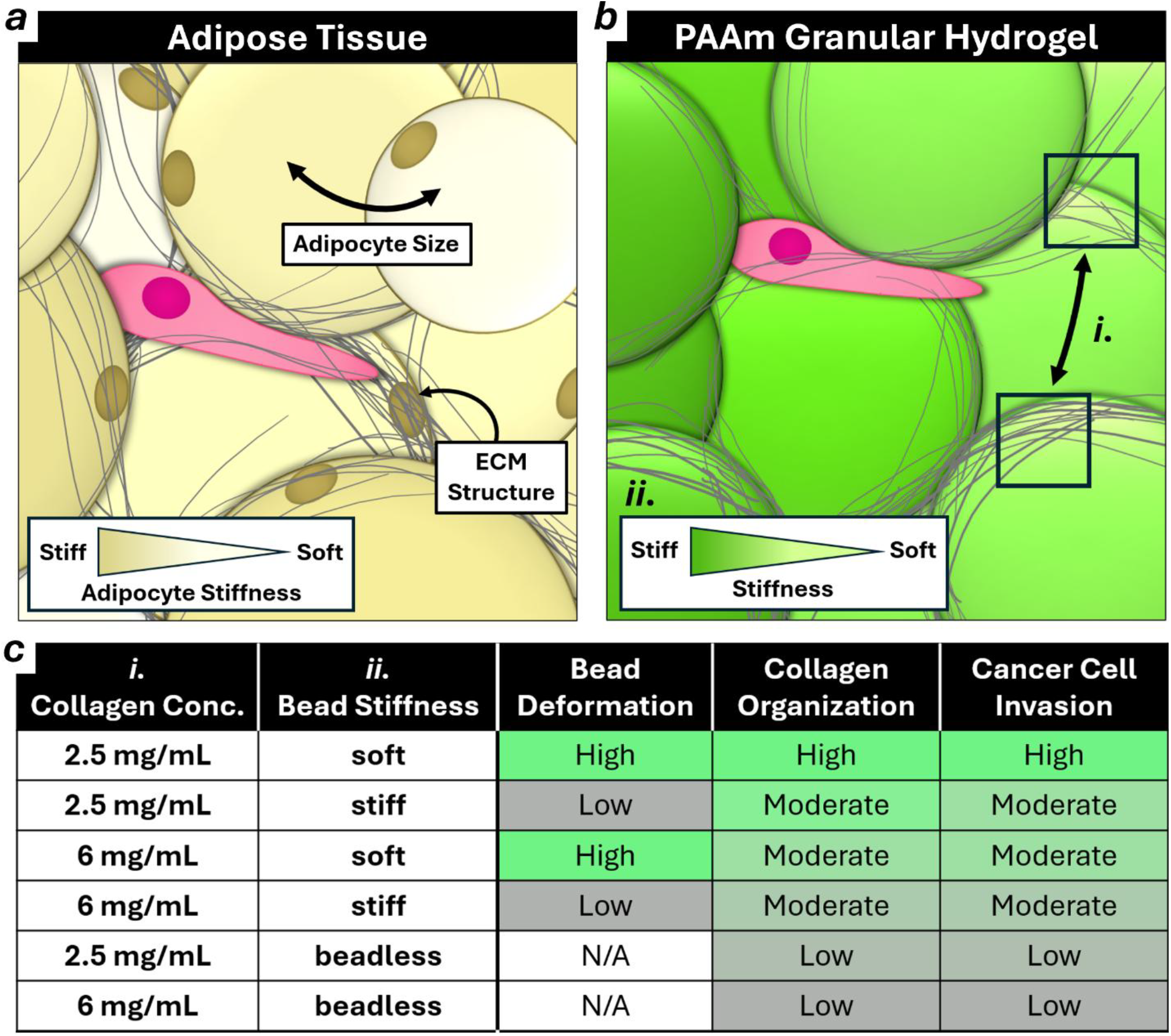
PAAm bead granular hydrogels recapitulate adipose tissue structure and alter cancer cell invasion in a stiffness-dependent manner. **(*a*)** Schematic depicting a motile cell migrating through adipose and the mechanically heterogenous stromal components which compose local tissue structure. **(*b*)** Schematic depicting a motile cell migrating through a PAAm bead granular hydrogel, mimetic of the highlighted adipose tissue properties via tunable ECM (*i*.) and bead (*ii*.) mechanics. **(*c*)** Table describing the hydrogel properties, corresponding to (*b*), which impact cell invasion phenotype.

This theoretical model offers insight into the cell-scale mechanical regulation of invasive cancer cells by native stromal adipocytes, emphasizing the migration-facilitating alignment of collagen fibers parallel to bead (adipocyte) interfaces and the rate-limiting, stromal stiffness-dependent confinement of migrating cells. The model is also consistent with findings of many previous studies, which describe the morphological plasticity migratory cells adapt as they encounter physically restrictive stromal structures or anisotropic ECM substrates^70,86–88^, as well as the unique rate-limiting but force-generating properties of confined cell nuclei^68,69,72^. Future validation of the model could be achieved by perturbing the ability of migratory cells to form traction-generating cytoskeletal protrusions^88,89^, as well as disrupting the structural integrity of the nuclear envelope^90^. Moreover, additional intravital imaging of other breast cancer cells migrating through native murine mammary adipose^91,92^ is needed to confirm the broad relevance of our findings, with the added potential to inform clinical prognostic imaging techniques in the future.

Like all model systems, adipose-mimetic granular hydrogels present some biological and technical limitations. For example, we used consistently sized particles to limit the confounding effects of varied bead curvature in polydisperse systems^93^ that can affect collagen fibrillogenesis and migration independently^22,94^. As native adipocyte size is heterogeneous, future studies with mixtures of differently sized PAAm particles will gain further insights into how adipocyte size and its heterogeneity affect our findings. Interestingly, a recently published study employing a biphasic hydrogel system suggests that ovarian cancer invasion mode is regulated by particle size and volume fraction, further demonstrating the importance of adipose tissue structure in regulating cancer progression^38^. Another interesting modification of the current platform would be to functionalize the surface of PAAm beads with adhesion ligands that recapitulate adipocyte-ECM interactions. Additionally, our model lacks other relevant stromal cell populations, ECM components, and adipokines secreted by native adipocytes. Some granular hydrogels systems have been leveraged to study other cell types, and have, for example, demonstrated how cancer cell phenotype and matrix composition coordinate adipose-derived stromal cell (ADSC) transition into cancer associated fibroblasts (CAFs)^37^. As our platform lends itself to incorporate these factors directly into the collagen matrix or the culture media, the role of other cell types and or biochemical factors could be further explored in the future. Lastly, due to the light-scattering properties of PAAm, imaging resolution through these granular hydrogels deteriorates as acquisition depth increases, restricting the hydrogel size as well as cell culture period over which imaging is possible. Here, we cast granular hydrogels into micron-sized PDMS molds to overcome these limitations, while excluding artifacts caused by limited transport of oxygen and other canonical morphogens^95^. Regardless, insights gathered from our system could inform future studies into the mechanical regulation of solid tumors arising in adipose-rich stroma (skin, ovarian, prostate, etc.)^96^, and have the potential to make a clinical impact by informing prognostic and diagnostic criteria across cancer types. Ultimately, our analyses highlight the impact of adipocyte and ECM mechanics on tumor cell invasion and demonstrate the need to further elucidate the mechanisms by which adipose stroma regulates breast cancer invasion.

## Resource availability

All source data and code to support experimental findings will be provided upon reasonable request to the corresponding author.

## Acknowledgments

We thank all members of the Fischbach Lab for valuable discussion and input during the preparation of this manuscript; Rebecca Williams for assistance with confocal microscopy; Ben Hopkins at WCM for help with generating *in vivo* tumors invading adipose tissue; the Cornell Center for Animal Resources and Education (CARE) for assistance with animal studies; the Cornell Animal Health Diagnostic Core for assistance with paraffin embedding and sectioning; and the Cornell Statistical Consulting Unit for assistance with statistical analysis. We acknowledge financial support from the NSF DGE2139899 (B.K.K), NCI F31CA278410 (G.F.B.), NCI R01CA259195 (C.F.) and R01CA276392 (C.F.), the Center on the Physics of Cancer Metabolism NCI 1U54CA210184 (C.F.), and the Cornell Engineering Learning Initiatives program (M.I.P.). We also acknowledge support by a Rosalind Franklin fellowship from the Max Planck Centre for Physics in Medicine (C.F.), and the Cornell Biotechnology Resource Center Imaging Facility funded by NYSTEM C029155, NIH S10OD018516, and NIH S10RR025502.

## Author contributions

B.K.K., G.F.B., C.S.O, and C.F. conceived and designed experiments. B.K.K. performed all experiments unless otherwise noted. B.K.K., G.F.B, and B.E.S. conducted initial system troubleshooting and protocol development. A.B. and G.F.B collected murine adipose tissue for histological analysis and primary adipocyte isolation, established adipocyte and bead size distributions, and worked with A.B. to survey bead/adipocyte stiffness using atomic force microscopy. S.G., J.G., and R.G. conceived, designed, and fabricated the adipocyte-mimetic polyacrylamide beads. D.W. computationally evaluated granular hydrogel packing fraction, collagen fiber eccentricity, and adipocyte diameter accuracy, and aided B.K.K. in confocal image processing. B.K.K, D.W., and C.S.O evaluated collagen fiber coherency data to define the network connectivity metric. Y.Z. calculated and analyzed breast cancer cell diffusion constants. C.E. and N.N. conceptualized and performed all intravital mammary fat pad imaging experiments, and B.K.K. processed the image data. D.W. and C.S.O. employed DEM modeling to predict cancer cell invasion dependent on adipocyte stiffness. B.K.K. prepared all figures and schematics. B.K.K, G.F.B., and C.F. wrote the manuscript with input from all authors.

## Supplemental data

Figures S1-S5

