## Supplemental Figures for "Adipose-mimetic granular hydrogels uncover biophysical cues driving breast cancer invasion"

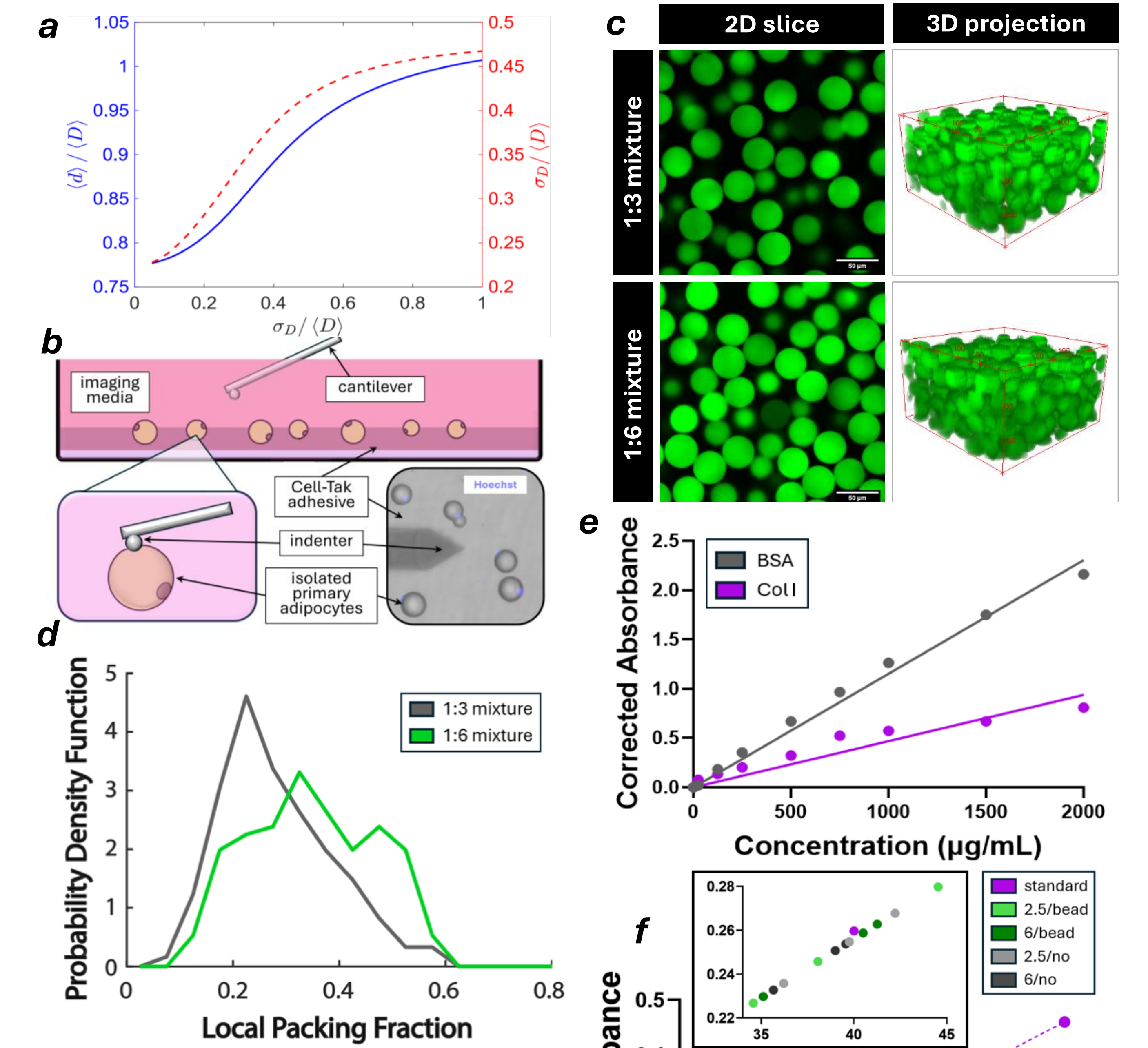


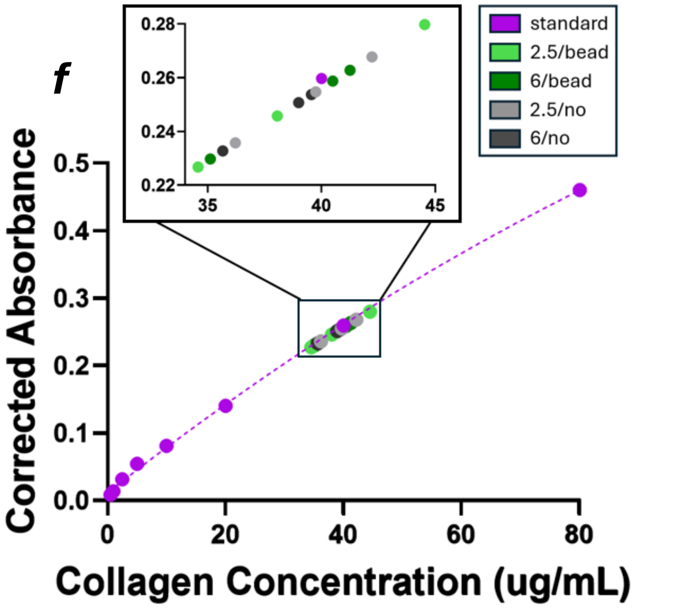


**g**

| Sample | Interpolated Conc. (ug/mL) | Dilution Factor | Original Conc. (mg/mL) |
|---|---|---|---|
| 2.5/bead | 39.00 | 58.8 | 2.29 |
| 6/bead | 38.94 | 142.9 | 5.56 |
| 2.5/no | 39.37 | 58.8 | 2.32 |
| 6/no | 38.06 | 142.9 | 5.44 |

**Supplementary Figure 1: Granular hydrogel system troubleshooting and characterization.**
***(a)*** Plot depicting the monotonic relationship between mean estimated ($\langle d \rangle$) and true ($\langle D \rangle$) adipocyte diameters, as depicted by proportional standard deviations ($\sigma$). ***(b)*** Schematic of atomic force microscopy adipocyte nanoindentation configuration including a representative image with Hoechst-labeled adipocytes. ***(c)*** Confocal micrographs depicting bead packing ratios of 1:3 and 1:6, scale = 50 µm. ***(d)*** Analysis of the local packing fraction of PAAm beads in hydrogel mixtures varying in collagen-to-bead ratio. ***(e)*** BCA assay standard curve comparisons of bovine serum albumin and collagen I. ***(f)*** BCA assay analysis of hydrogel collagen concentrations interpolated from a collagen I standard curve. ***(g)*** Mean interpolated collagen concentration values for beadless and granular hydrogels, accounting for initial target dilution of samples to 40 µg/mL.

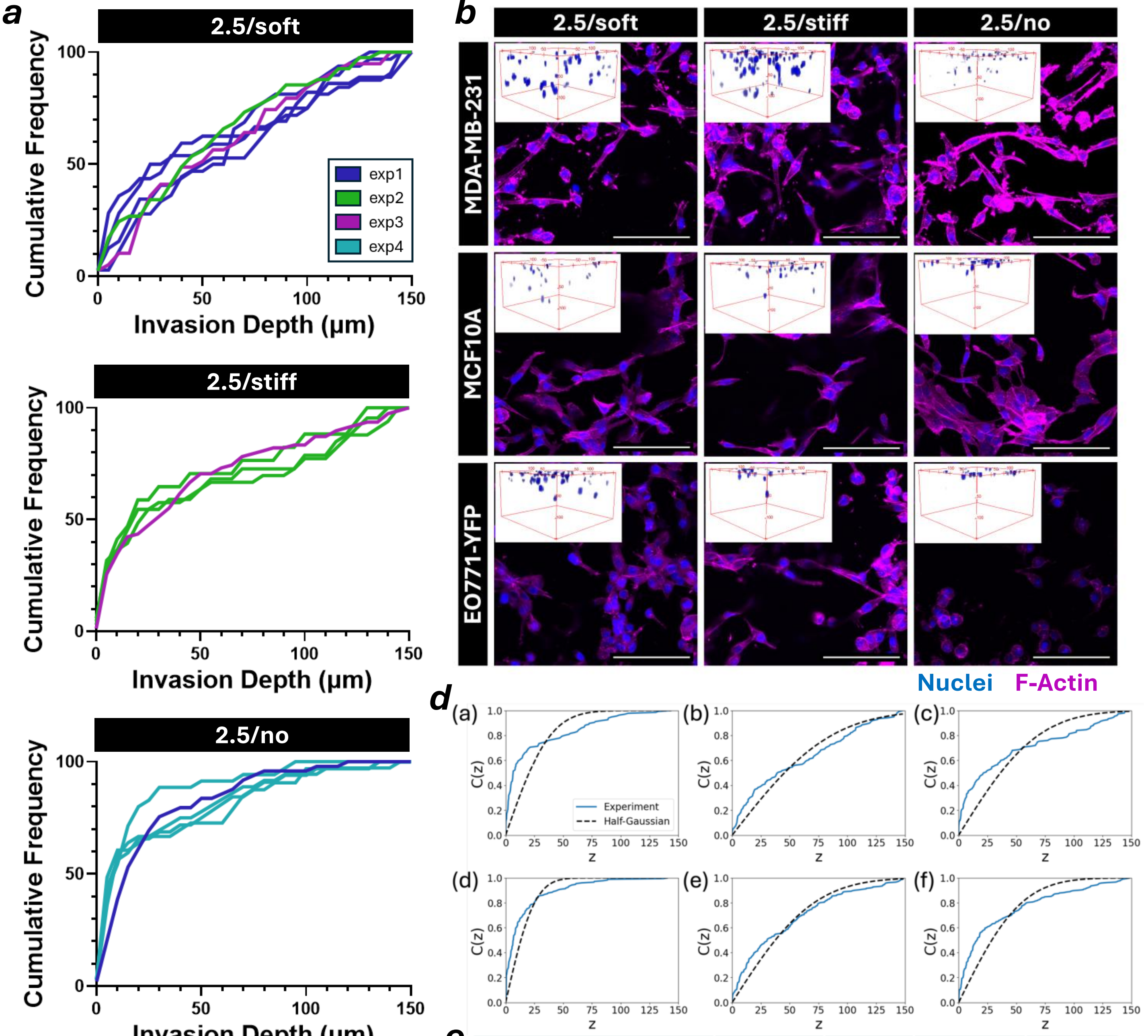


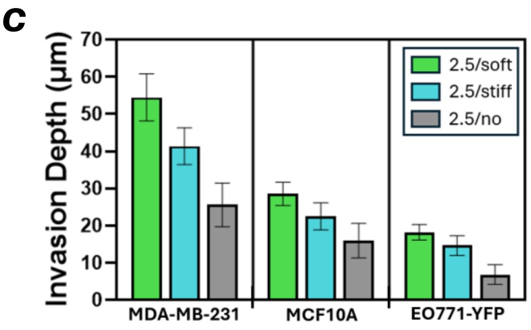


**e**

| | condition | sigma | D | std(D) | $R^2$ |
|---|---|---|---|---|---|
| 2.5 mg/mL | (a) beadless | 24.1 | 12.7 | 2.8 | 0.52 |
| | (b) soft bead | 53.7 | 62.9 | 8.2 | 0.93 |
| | (c) stiff bead | 42.9 | 40.2 | 6.4 | 0.78 |
| 6 mg/mL | (d) beadless | 15.6 | 5.3 | 1.2 | 0.69 |
| | (e) soft bead | 44.3 | 42.7 | 4.4 | 0.93 |
| | (f) stiff bead | 34.0 | 25.3 | 3.3 | 0.78 |

**Supplementary Figure 2: Validation of mammary cell invasion through adipose-mimetic hydrogels.**
(***a***) Plots evaluating the interexperimental variability of MDA-MB-231 cells invading through hydrogels composed of 2.5 mg/mL collagen and soft, stiff, or no PAAm beads. (***b***) 150 µm maximum projections and 3D reconstructions of MDA-MB-231, MCF10A, and EO771 mammary cell lines invading hydrogels comprised of 2.5 mg/mL collagen and soft, stiff, or no PAAm beads after 36 hours, scale = 100 µm. (***c***) Mean +/- SEM invasion depth of various mammary cell lines through granular hydrogels varying in mechanical properties. (***d***) Plots depicting the invasion distributions of cells migrating through (a) 2.5 mg/mL collagen beadless, (b) 2.5 mg/mL collagen soft bead, (c) 2.5 mg/mL collagen stiff bead, (d) 6 mg/mL collagen beadless, (e) 6 mg/mL collagen soft bead, and (f) 6 mg/mL collagen stiff bead hydrogels compared to a half-gaussian distribution, and (***e***) corresponding coefficients.

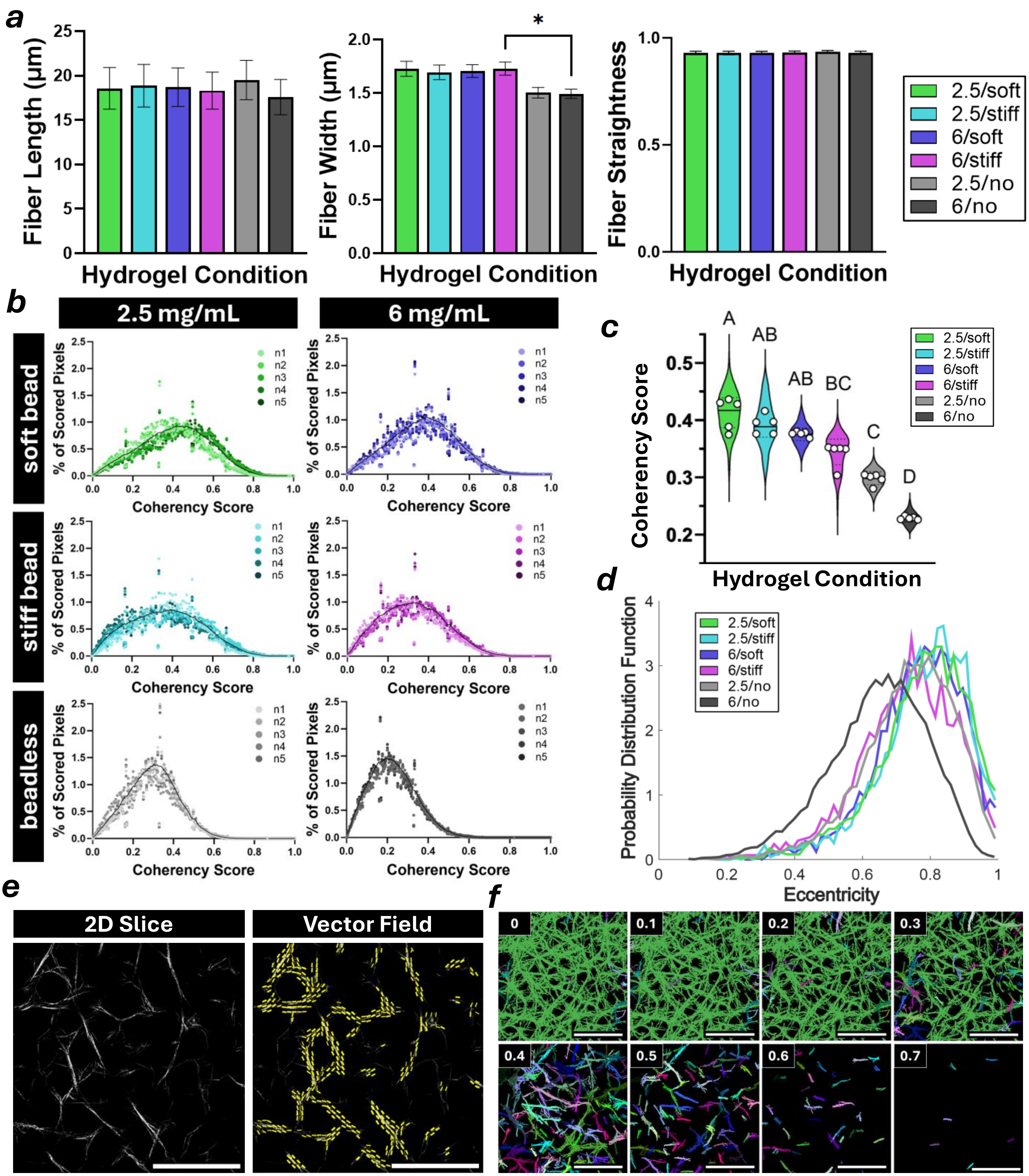


**Supplementary Figure 3: Validation of collagen I fiber characteristics in adipose-mimetic hydrogels.** (***a***) Plots depicting the mean length, width, and straightness of collagen fibers across various hydrogel types, as evaluated by the MATLAB plugin CurveAlign. Error bars represent SEM. (***b***) Plots evaluating the interexperimental variability of type I collagen fiber coherency score measurements. **(c)** Violin plots demonstrating average collagen fiber coherency scores depending on hydrogel condition. (***d***) Collagen fiber eccentricity analysis as a relative metric of fiber curvature, corresponding to (***e***) a representative reflectance micrograph overlaid with the computed vector field,  scale = 100 μm. (***f***) Collagen fiber network thresholding according to increasingly stringent coherency scores, used to determine network interconnectivity, scale = 100 μm.

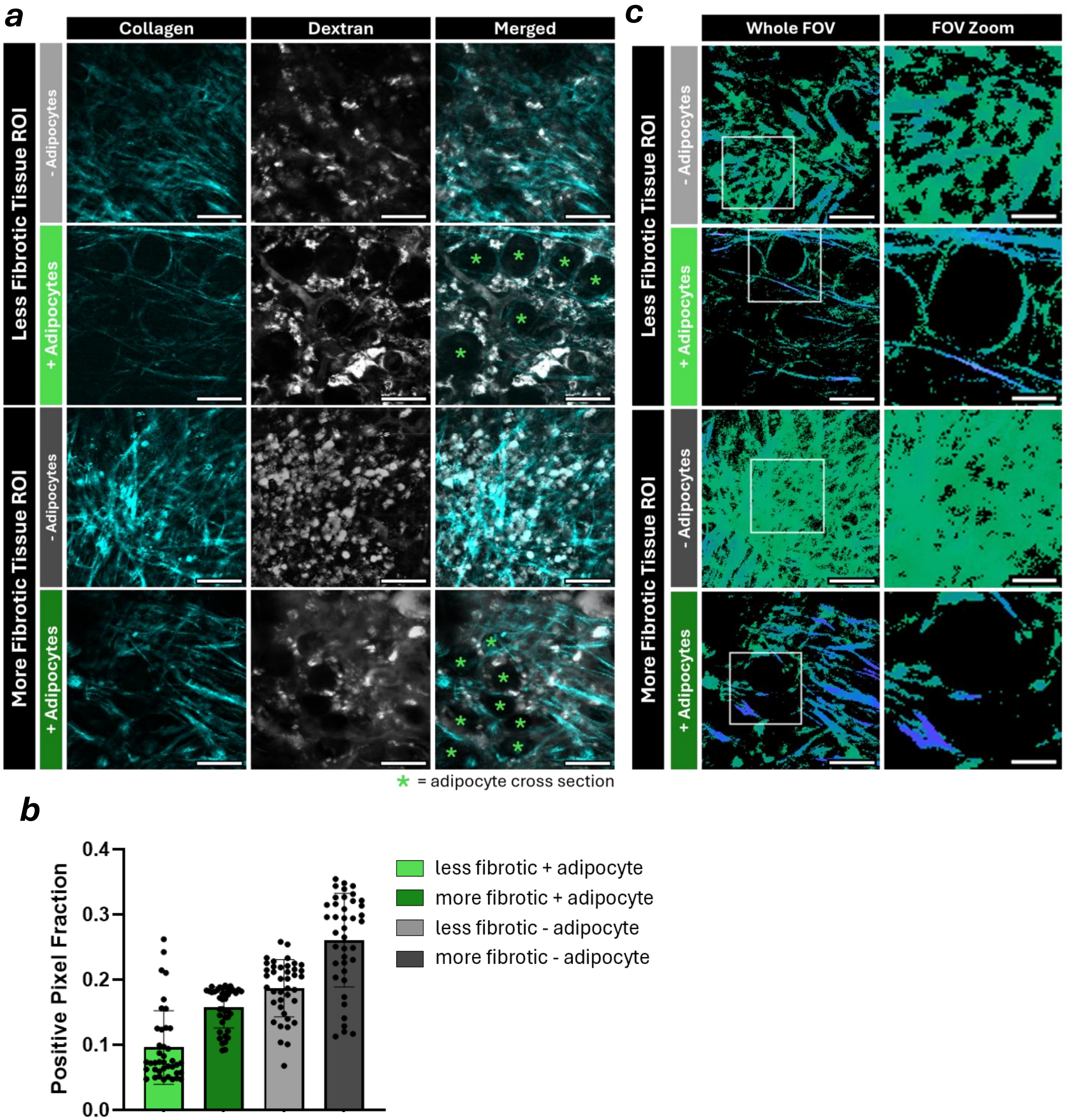


**Supplementary Figure 4: Validation of cancerous mammary fat pad collagen characteristics *in vivo*.** (***a***) An expansion of Fig. 4f, 10 µm maximum projection SHG (collagen) and fluorescent (dextran) micrographs capturing varying levels of fibrosis and adipocyte packing in live murine mammary tissue, scale = 25 µm. (***b***) Corresponding to Fig. 4f, quantification of local degree of tissue fibrosis via collagen channel thresholding. (***c***) An expansion of Fig. 4g, coherency color survey analysis of 10 µm maximum projection SHG micrographs depicting collagen fiber structure in living murine mammary fat pads, with whole (left, scale = 25 µm) and zoomed (right, scale = 10 µm) fields of view.

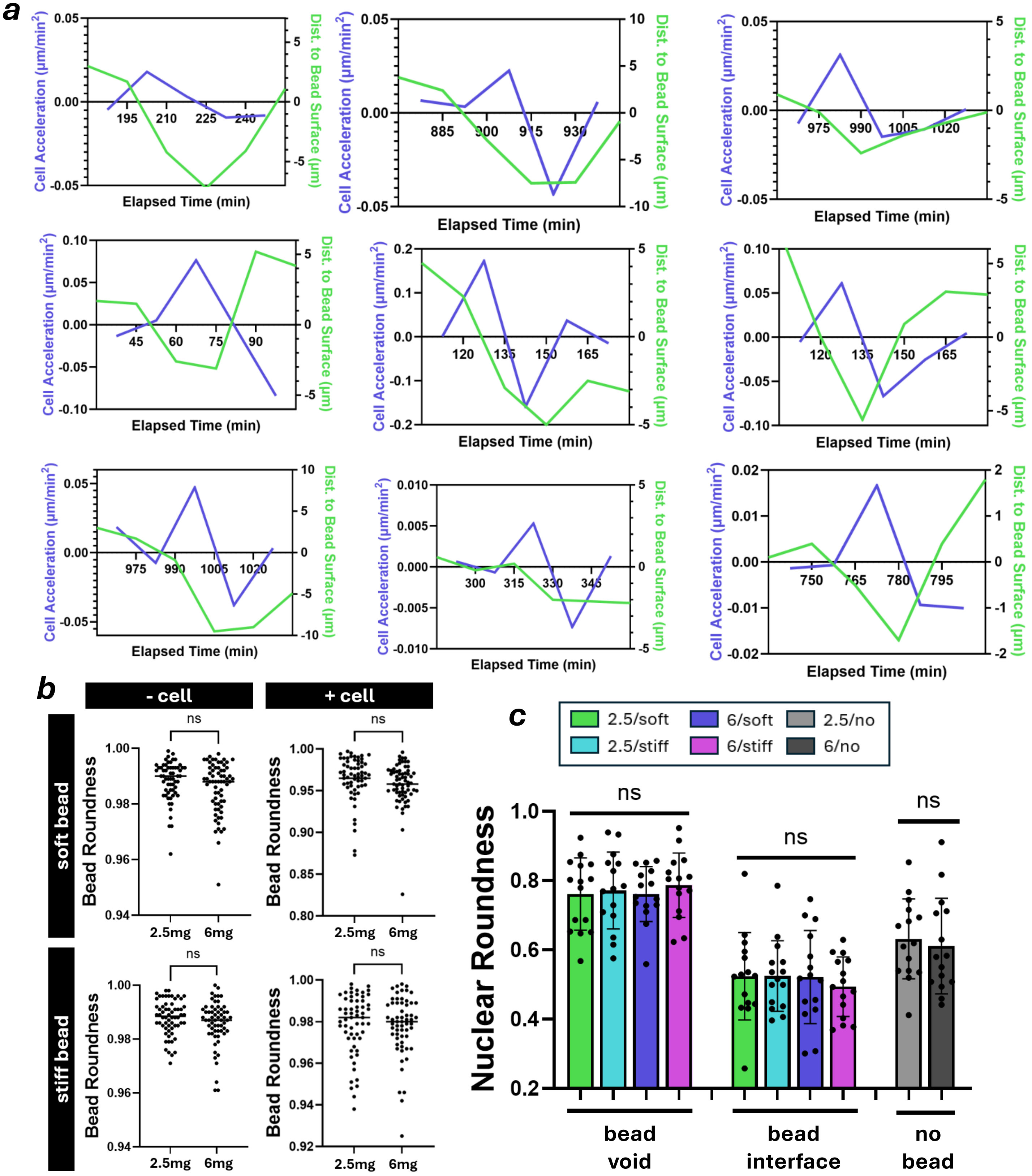


**Supplementary Figure 5: Characterization and validation of bead and nuclear properties during cancer cell migration.** (***a***) Plots depicting several distinct moments of nuclear acceleration and deceleration coordinated by confined bead interfaces. (***b***) Plots demonstrating the shape factor consistency (roundness) of PAAm beads embedded within 2.5 or 6 mg/mL collagen hydrogels, with or without contact by migrating MDA-MB-231 nuclei. (***c***) A comparison of migratory cell nuclei shape factors (roundness) between various hydrogel conditions, where nuclei are binned based on localization within unconfined void regions or confined bead interface regions.
